# Impact of prior and projected climate change on US Lyme disease incidence

**DOI:** 10.1101/2020.01.31.929380

**Authors:** Lisa I. Couper, Andrew J. MacDonald, Erin A. Mordecai

**Affiliations:** Department of Biology, Stanford University, Stanford, California; Earth Research Institute, University of California, Santa Barbara, California; Bren School of Environmental Science and Management, University of California, Santa Barbara, California

**Author notes:** **Corresponding author:** Lisa Couper, 327 Campus Drive, Stanford, CA 94305. The authors declare they have no actual or potential competing financial interests*.

**Keywords:** Lyme disease, climate change, *Ixodes scapularis*, *Ixodes pacificus*, least squares dummy variables, disease projections

## Abstract

Lyme disease is the most common vector-borne disease in temperate zones and a growing public health threat in the United States (US). The life cycles of the tick vectors and spirochete pathogen are highly sensitive to climate, but determining the impact of climate change on Lyme disease burden has been challenging due to the complex ecology of the disease and the presence of multiple, interacting drivers of transmission. Here we incorporated 18 years of annual, county-level Lyme disease case data in a panel data statistical model to investigate prior effects of climate variation on disease incidence while controlling for other putative drivers. We then used these climate-disease relationships to project Lyme disease cases using CMIP5 global climate models and two potential climate scenarios (RCP4.5 and RCP8.5). We find that interannual variation in Lyme disease incidence is associated with climate variation in all US regions encompassing the range of the primary vector species. In all regions, the climate predictors explained less of the variation in Lyme disease incidence than unobserved county-level heterogeneity, but the strongest climate-disease association detected was between warming annual temperatures and increasing incidence in the Northeast. Lyme disease projections indicate that cases in the Northeast will increase significantly by 2050 (23,619 ± 21,607 additional cases), but only under RCP8.5, and with large uncertainty around this projected increase. Significant case changes are not projected for any other region under either climate scenario. The results demonstrate a regionally variable and nuanced relationship between climate change and Lyme disease, indicating possible nonlinear responses of vector ticks and transmission dynamics to projected climate change. Moreover, our results highlight the need for improved preparedness and public health interventions in endemic regions to minimize the impact of further climate change-induced increases in Lyme disease burden.

## Introduction

Arthropod-transmitted pathogens pose a severe and growing threat to global public health (World Health Organization 2014). Because vector life cycles and disease transmission are highly sensitive to abiotic conditions (Mattingly 1969, Sonenshine and Roe 2013), climate change is expected to alter the magnitude and geographic distribution of vector-borne diseases (Kilpatrick and Randolph 2012, World Health Organization 2014). Climatic changes, in particular warming temperatures, have already facilitated expansion of several vector species (e.g., Purse et al. 2005, González et al. 2010, Roiz et al. 2011, Clow et al. 2017a), and have been associated with increased vector-borne disease incidence (e.g., Loevinsohn 1994, Subak 2003, Hii et al. 2009). Identifying areas of high risk for current and future vector-borne disease transmission under climate change is critical for mitigating disease burden. However, the presence of interacting drivers of disease transmission such as land use change and globalization, and the complex ecology of vector-borne diseases make the effort to measure and predict effects of climate on vector-borne disease incidence challenging (Rogers and Randolph 2006, Tabachnick 2010, Mills et al. 2010, Ostfeld and Brunner 2015, Lafferty and Mordecai 2016).

This challenge is particularly apparent in the case of Lyme disease, the most common vector-borne disease in temperate zones (Kurtenbach et al. 2006, Rizzoli et al. 2011, Rosenberg et al. 2018), because transmission depends on a complex sequence of biotic interactions between vector and numerous host species that may respond differently to environmental change (Ostfeld 1997). In the United States (US), Lyme disease is caused by the bacteria *Borrelia burgdorferi*, and is vectored by two tick species: *Ixodes scapularis* in the eastern and midwestern US and *Ixodes pacificus* in the western US. After hatching from eggs, both tick species have three developmental stages—larva, nymph, and adult—during which they take a single blood meal from a wide range of vertebrate hosts before transitioning to the next developmental stage or reproducing (Sonenshine and Roe 2013). This life cycle takes 2-3 years to complete, 95% of which is spent at or below the ground surface in diapause, seeking a host, digesting a blood meal, or molting (i.e., off the host) (Sonenshine and Roe 2013, Ostfeld and Brunner 2015).

Given their long life spans, ectothermic physiology, and high degree of interaction with the physical environment, tick life cycles are sensitive to changes in climate and weather conditions (Sonenshine and Roe 2013). Prior research has demonstrated that temperature and moisture strongly influence tick mortality, development, and host-seeking abilities (reviewed in Ostfeld and Brunner 2015, Ogden and Lindsay 2016). In particular, both low and high temperatures decrease *I. scapularis* and *I. pacificus* survival and host-seeking activity (Lindsay et al. 1995, Vandyk et al. 1996, Padgett and Lane 2001). Further, cool temperatures prolong tick development and increase generation times, leading to greater proportional mortality before reproduction (Peavey and Lane 1996, Ogden et al. 2004, 2006). Rainfall and moisture availability also influence host-seeking activity in nonlinear ways. Low humidity exposure substantially increases tick mortality and inhibits host-seeking activity (Stafford 1994, Lane et al. 1995, Vail and Smith 1998, Schulze et al. 2001, Rodgers et al. 2007, Nieto et al. 2010, Ginsberg et al. 2017, MacDonald et al. 2019b). To avoid desiccating conditions, Ixodid ticks often modify their questing behavior to remain closer to the moist vegetative surface, or return frequently to rehydrate, both of which decrease the probability of obtaining a blood meal and thereby limiting survival and reproduction (Randolph and Storey 1999, Prusinski et al. 2006, Sonenshine and Roe 2013, Arsnoe et al. 2015, McClure and Diuk-Wasser 2019). However, heavy rainfall may also directly impede tick host-seeking (Randolph 1997). Given these physiological relationships, temperature and precipitation are important predictors of these tick species’ latitudinal and altitudinal range limits (McEnroe 1977, Estrada-Peña 2002, Brownstein et al. 2003, Ogden et al. 2005, Leighton et al. 2012, Berger et al. 2014, Eisen et al. 2016, Hahn et al. 2016), and northward range expansion of *I. scapularis* has been associated with warming temperature (Ogden et al. 2014b, Clow et al. 2017b, 2017a).

Yet despite well-known physiological relationships between specific climate variables and aspects of tick biology, and strong evidence of relationships between climate and tick range limits, it remains unclear how these effects translate into Lyme disease incidence - the outcome of interest to public health - and how broadly they apply across biogeographically distinct US regions. However, associations between climate and Lyme disease incidence are difficult to measure given the influence of many non-climate factors such as changing physician awareness, host movement, and human behavior (Morshed et al. 2006, Randolph 2010, Ostfeld and Brunner 2015, Kilpatrick et al. 2017, Scott and Scott 2018). A handful of prior studies have attempted to isolate the effect of climate on incidence, but have been limited in geographic or temporal scope, and/or not controlled for confounding drivers of incidence, leading to conflicting results about the role of climate change on transmission (Subak 2003, McCabe and Bunnell 2004, Schauber et al. 2005, Burtis et al. 2016, Dumic and Severnini 2018). As a result, our ability to predict effects of future climate change on Lyme disease incidence remains limited.

Here, we leverage an 18-year county-level Lyme disease case reporting dataset and explicitly control for other drivers of disease burden to ask: How has interannual variation in climate conditions contributed to past changes in Lyme disease incidence across distinct US regions? We include climate variables capturing changes in temperature and precipitation conditions and investigate how relationships between climate and Lyme disease outcomes vary across different regions of the US (i.e., the Northeast, Midwest, Southeast, Southwest, Pacific Southwest, and Pacific). We hypothesize that: a) warmer temperatures in northern regions and b) spring precipitation in all regions promote tick survival and therefore increase Lyme disease incidence, while c) hot, dry conditions during the questing period decrease tick host-seeking activity, survival and disease incidence. To avoid drawing spurious conclusions about the effects of climate, we analyze the effects of other known and potential drivers of disease incidence such as changing forest cover, public awareness of tick-borne disease, and health-seeking behavior, and use a statistical approach that explicitly accounts for unobserved heterogeneity in disease incidence between counties and years. We then use these modeled, regionally-specific relationships between climate and Lyme disease burden to investigate projected changes in US Lyme disease incidence under future climate scenarios. We report the projected change in Lyme disease incidence for individual US regions in 2040 – 2050 and 2090 – 2100 relative to hindcasted 2010 – 2020 levels under two potential climate scenarios: RCP8.5, which reflects the upper range of the literature on emissions, and RCP4.5, which reflects a moderate mitigation scenario (Hayhoe et al. 2017).

## Materials and Methods

### Lyme disease case data

We obtained annual, county-level reports of Lyme disease cases spanning from 2000 to 2017 from the US Centers for Disease Control and Prevention (CDC) (see Supporting Information). These disease case data provide the most spatially-resolved, publicly available surveillance data in the US. Raw case counts were converted to incidence using annual county population sizes from the US Census Bureau (USCB) and were expressed in cases per 100,000 people.

### Climate data

An overwhelming number of climate variables, such as the mean, range, and maximum or minimum temperature or precipitation at different time scales, could conceivably affect Lyme disease transmission. To reduce the probability of identifying significant but spurious relationships between climate and incidence, we limited the variables considered here to: average winter temperature lagged 1.5 years; average spring precipitation; the number of hot, dry days in May – July (the nymphal tick questing period); cumulative average temperature; total annual precipitation; daily temperature variability; and daily precipitation variability (Table 1). These variables have either been previously associated with variation in Lyme disease incidence, tick range limits or abundance, or, in the case of daily temperature and precipitation variability, are grounded in physiological relationships between climate and tick life history but have not been previously tested. In particular, interannual variation in Lyme disease incidence in endemic regions has been positively associated with lagged average winter temperature (Subak 2003), average spring precipitation (McCabe and Bunnell 2004), and negatively associated with the number of hot, dry days in May – July (Burtis et al. 2016). A measure of cumulative annual temperature (degree days > 0°C) has been associated with *I. scapularis* population establishment and abundance (Jones and Kitron 2000, Ogden et al. 2004, 2006, Clow et al. 2017b), and cumulative annual precipitation has been associated with larval tick abundance (Jones and Kitron 2000). Frequent variation in temperature can decrease tick survival due to the energetic costs of adapting to changing conditions (Gigon 1985, Herrmann and Gern 2013), thus daily temperature and precipitation variability were included here to explore whether this effect scaled to affect transmission risk. Details about how these variables were calculated and further justification for their biological relevance are listed in Table 1.

**Table 1.** Climate variables considered for models of disease incidence by region, along with descriptions and justification of their relevance to disease transmission.

| Climate Variable | Description | Biological Relevance |
| --- | --- | --- |
| Lagged winter temperature | Average monthly temperatures for Dec - Feb 1.5 years prior. Identified by Subak 2003 as significantly positively correlated with Lyme disease incidence in highly endemic areas. | Colder winter temperatures are associated with reduced host-seeking abilities of the adult tick (Duffy and Campbell 1994, Clark 1995, Carroll and Kramer 2003) and reduced abundance of the white-footed mouse, a highly competent reservoir host (Wolff 1996). |
| Spring precipitation | Average precipitation in May and June. Identified by McCabe and Bunnell 2004 as significantly positively correlated with Lyme disease incidence in highly endemic areas. | Greater precipitation during the late spring and early summer increases the moisture of the leaf litter, providing conditions which promote the survival and questing activity of the nymphal life stage (Knülle and Rudolph 1982, Berger et al. 2014). |
| Hot, dry days | The number of days with temperature > 25°C and precipitation = 0 during May – July (or May – June for counties with *Ixodes pacificus*). Identified by Burtis et al. 2016 as significantly negatively correlated with Lyme disease incidence in highly endemic areas. | Hot, dry conditions are associated with decreased questing activity and questing height of ticks (Randolph and Storey 1999, Schulze et al. 2001), reducing the likelihood of attachment to humans (Arsnoe et al. 2015). The May - July, and May - June, time periods capture the peak nymphal questing periods for *I. scapularis* and *I. pacificus*, respectively (Eisen et al. 2016). |
| Cumulative average temperature | The sum of average daily temperatures (°F) over the entire year | Cumulative temperature appears to control most developmental stages of *I. scapularis* (Lindsay et al. 1995, Rand et al. 2004). Lower cumulative temperature is associated with longer development periods and/or higher tick mortality (McEnroe 1977, Estrada-Peña 2002, Brownstein et al. 2003, Ogden et al. 2004, Leighton et al. 2012). |
| Total annual precipitation | The sum of total daily precipitation (mm) over the entire year | Greater precipitation increases the moisture of the leaf litter, providing conditions which favor tick survival and questing activity (Knülle and Rudolph 1982, Jones and Kitron 2000, Berger et al. 2014a). |
| Daily temperature variability | The variance in average daily temperatures (°F) over the entire year | Frequent temperature variation can decrease tick survival, even beyond that of constant cold exposure, due to energetic costs associated with adapting to changing temperatures (Gigon 1985, Hermann and Gern, 2013); however, effects will vary based on the average temperature of the region. |
| Daily precipitation variability | The variance in total daily precipitation (mm) over the entire year | Both drought and heavy rainfall are associated with decreased tick questing activity and survival (Randolph 1997, Jones and Kitron 2000, Perret et al. 2004). Variation in precipitation, as opposed to consistent rainfall supplying favorable high relative humidity conditions, may thus be detrimental for tick survival, but will depend on the average precipitation of the region and the magnitude of variation. |

For past climate conditions, we obtained daily, county-level average temperature and total precipitation data from the National Oceanic and Atmospheric Administration (NOAA) weather stations accessed via the CDC’s Wide-ranging Online Data for Epidemiological Research (WONDER) database. To estimate future climate variables, we used NASA Goddard Institute for Space Studies CMIP5 data on modeled temperature and precipitation (Schmidt et al. 2014). Specifically, we obtained estimates of daily near-surface air temperature and precipitation through 2100 under the upper climate change scenario (RCP8.5) and a moderate climate change scenario (RCP4.5) (van Vuuren et al. 2011, Taylor et al. 2012). These climate scenarios are relatively similar in the radiative forcing levels assumed through 2050 but diverge substantially in the latter half of the century. Climate estimates from these two scenarios are provided at a 2° x 2.5° resolution; values were then ascribed to counties based on county latitude and longitude (see Figure S1). Mean values for hindcasted and projected climate variables for each region are listed in Table S1.

### Awareness data

We controlled for variation in public awareness of ticks and Lyme disease using data from Google trends on the frequency of “ticks” as a search term. We obtained data on “ticks” search frequency, normalized for a given location and year, for 2004 (the first year the data were available) to 2017. We also initially used “tick bite”, and “Lyme disease” as search terms, but found that these generated nearly identical coefficient estimates, thus we proceeded to use only the “ticks” search term as a predictor. Search frequency data were aggregated at the designated market area (DMA), the smallest spatial scale available. Search frequency values for a given DMA, which contained an average of 14 counties, were applied equally to all counties therein. We used a 1-year lagged version of the tick search variable, as awareness of tick-borne disease is likely endogenous to incidence (i.e., higher Lyme disease incidence likely contributes to higher tick search frequency and awareness) and using predetermined values reduces endogeneity concerns (Bascle 2008).

### Health-seeking behavior data

We explicitly controlled for variation in health-seeking behavior, previously posited as a driver of Lyme disease reporting (Armstrong et al. 2001, Wilking and Stark 2014) by including health insurance coverage and poverty as potential predictors. Given the logistical and financial challenges in obtaining a Lyme disease diagnosis and treatment (Johnson et al. 2011, Adrion et al. 2015), access to health care services may play a role in whether a Lyme disease case is identified and reported. We obtained data on health insurance coverage, defined as the percent of county residents with any form of health insurance coverage in a given year, for 2005 to 2017 from USCB’s Small Area Health Insurance Estimates (SAHIE) program. We obtained data on poverty, defined as the percent of county residents living in poverty in a given year, for 2000 to 2017 from the USCB.

### Land cover data

We included two land cover variables putatively associated with higher tick-borne disease risk: the percent forest in a given county and year, and the percent mixed development (Brownstein et al. 2005b, Dister and Fish 1997, Frank et al. 1998, Glass et al. 1995, Killilea et al. 2008, MacDonald et al. 2019a). We calculated these variables using 30-m resolution land cover data from the US Geological Survey (USGS) National Land Cover Database (NLCD) (Yang et al. 2018). Percent forest included any deciduous, evergreen, or mixed forest. Mixed development was defined as areas with a mixture of constructed materials and vegetation, including lawn grasses, parks, golf courses, and vegetation planted in developed settings. We calculated county-level values of these land cover variables for 2001, 2004, 2006, 2008, 2011, 2013, and 2016 as these are the only years the NLCD dataset is currently available.

To estimate future land cover variables, we used land cover projections generated by the USGS Earth Resources Observation and Science Center (EROS) using the IPCC Special Report on Emissions Scenarios (SRES) (Sohl et al. 2014). Although newer socioeconomic pathways have recently been developed (i.e., the “Shared Socioeconomic Pathways”), these scenarios have not yet been incorporated into US land cover projections (Sohl 2019). We used modeled land cover data under SRES B1, which reflects lower urban development, to align with the moderate climate change scenario (RCP4.5), and SRES A1B, which reflects higher urban development and conversion of natural lands, to align with the upper climate change scenario (RCP8.5) (Nakicenovic et al. 2000, Rogelj et al. 2012, Sohl et al. 2014). Using these data, we again calculated annual, county-level values of percent forest cover and mixed development for 2040 – 2050 and 2090 – 2100. However, as the ‘mixed development’ land cover class was not included in the projected data, we instead used the ‘mechanically disturbed’ public or private land cover class (see Supporting Information).

### Regional divisions

Given the large variation in climatic conditions across the US, as well as variation in ecological dynamics of tick-borne diseases such as tick species identity, tick densities, tick questing behavior, and host community composition (Eisen et al. 2016, Kilpatrick et al. 2017, Ostfeld 1997, Salkeld and Lane 2010), we examined regional differences in climate-disease relationships. We used the US Fish & Wildlife Service regional boundaries to divide the US into the following seven regions for analysis: Northeast, Midwest, Mountain Prairie, Pacific, Pacific Southwest, Southwest, and Southeast (Figure 1). These regional divisions were selected as they roughly correspond to genetic structuring of *I. scapularis* and *I. pacificus* (Kain et al. 1997, 1999, Humphrey et al. 2010) and are likely distinct in environmental conditions and resources (Ricketts et al. 1999, Smith et al. 2018). These regional divisions are also similar to the nine ‘climatically consistent’ regions within the contiguous US identified by NOAA (Karl and Kloss 1984) but preserve larger regions in the South and Midwest to obtain higher power in the analysis. Further, each region contains only one vector species: *I. scapularis* in the Northeast, Midwest, Southeast, and Southwest, and *I. pacificus* in the Pacific and Pacific Southwest (Dennis et al. 1998). As neither species has an established presence in the Mountain Prairie, this region was removed from the analysis. Regional descriptions, including the population size (as of 2017), the number of counties, and the average climate conditions, are provided in Table S2.

**Figure 1.**
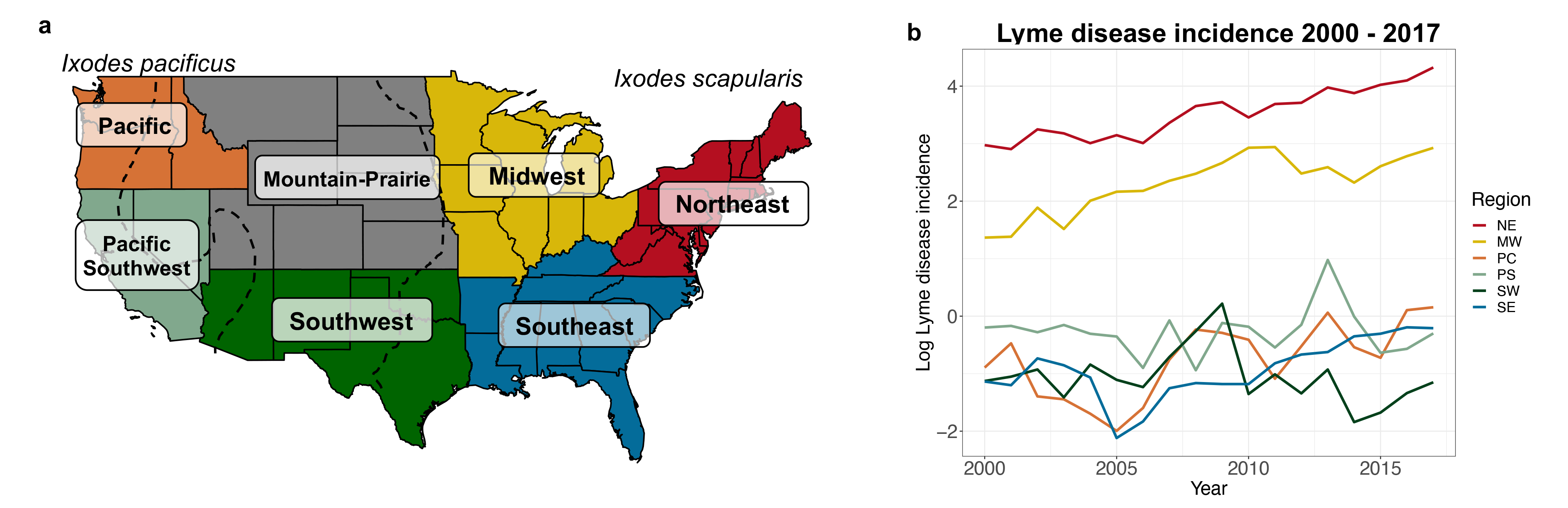
**a)** Regional boundaries designated by US Fish & Wildlife Service. These regions were used to analyze spatial variation in the effects of climate conditions on disease outcomes. Map recreated from: https://www.fws.gov/endangered/regions/index.html. Dashed black lines denote the approximate eastern boundary of *Ixodes pacificus* and western boundary of *Ixodes scapularis* based on distribution maps created by the CDC. **b)** Regional time series of log Lyme disease incidence (the number of cases per 100,000 people in the population) from 2000 – 2017. The Mountain Prairie region is not shown here as it was removed from the analysis due to low vector presence at the start of the analysis period.

### Statistical analysis

We used a least squares dummy variable (termed “fixed-effects” in econometrics) regression approach to estimate changes in Lyme disease incidence using repeated observations of the same groups (counties) from 2000 – 2017 (Larsen et al. 2019). This class of statistical approaches has been developed to isolate potential causal relationships in the absence of randomized experiments where such experiments are not feasible (Larsen et al. 2019, MacDonald and Mordecai 2019). We included ‘county’ and ‘year’ dummy variables to control for any unobserved heterogeneity that may influence reported Lyme disease incidence in a particular county across all years (e.g., geographic features, number of health care providers), or influence Lyme disease in all counties in a given year (e.g., changes in disease case definition), respectively. All counties (n = 2,232) for which there were complete data on Lyme disease cases, climate, and other predictors were included.

To account for regional variation in the predictors of tick-borne disease incidence (Wimberly et al. 2008, Raghavan et al. 2014), we ran separate models for each US region (see Methods: Regional divisions). We used stepwise variable selection, in which variables were added if they reduced model Akaike information criterion (AIC) by two or more, to identify the climate, land cover, and non-ecological predictors that best explained Lyme disease incidence in each region (Yamashita et al. 2007, Zhang 2016). We assessed the multicollinearity of these models by calculating the variance inflation factor (VIF). No predictors had VIF values greater than 10 after the stepwise variable selection procedure, thus we did not remove any variables from the final models due to high collinearity (Hair et al. 2014).

We accounted for spatial and temporal autocorrelation of model errors by using cluster-robust standard errors. This nonparametric approach accounts for arbitrary forms of autocorrelation within a defined “cluster” to avoid misleadingly small standard errors and test statistics (Cameron and Miller 2015). We specified clusters as US Agricultural Statistics Districts (ASDs), which contain on average 9.9 ± 5.2 counties. These districts contain contiguous counties grouped by similarities in soil type, terrain, and climate such that each district is more homogenous with respect to these characteristics than the state as a whole (USDA 2018). Accounting for spatial and temporal correlation in this way may help to account for ecological similarities between neighboring counties not captured in the climate and land cover predictors. Along these lines, ASDs have previously been used to account for spatial autocorrelation when investigating relationships between forest fragmentation and Lyme disease incidence at the county-level (MacDonald et al. 2019a). When reporting on the significance of a predictor, we use standard errors and p-values calculated using this correction. To ensure our results were robust to cluster specification, we repeated the model runs using county as the cluster unit (Table S3). All analyses were conducted in R version 3.6 (R Core Team 2017)

To capture any nonlinear relationships between climate predictors and Lyme disease incidence, we generated models using linear and quadratic versions of the climate variables as potential predictors. Specifically, we used the stepwise variable selection approach starting with linear and quadratic versions of each climate variable to determine the best fit model for each region. We compare model accuracy and the output of these models to those using only linear versions of climate predictors to assess the sensitivity of our results to the functional form of climate-disease relationships (see Methods: Model validation).

### Lyme disease projections

We projected Lyme disease incidence using the climate and land cover variables included in the best fit model for each region as well as a county dummy variable. Tick search frequency, poverty, and health insurance coverage were not included because annual, county-level projections for these variables are unavailable. Using these models, we obtained regional estimates for Lyme disease incidence under the upper and moderate climate change scenarios (RCP8.5 and RCP4.5) for 2040 – 2050 and 2090 – 2100. We calculated county-level changes in Lyme disease incidence by subtracting modeled incidence for 2010 – 2020 from projected incidence. Using modeled incidence for 2010 – 2020, rather than true case data for the years it was available, allowed for more direct comparisons between prior and projected cases because these estimates were made using the same climate and land cover data.

We converted projected Lyme disease incidence to cases under two differing assumptions about county population sizes. In the first calculation, we account for projected population growth by using county-level population projections under the Shared Socioeconomic Pathway “Middle of the Road” scenario (SSP2) as generated by Hauer 2019 (Samir and Lutz 2017). In the second, we assume that county population sizes remained the same as those in 2017, the last year of available county-level Lyme disease case reports. We focus our results and discussion on the projections made using population size projections, but compare results from these two approaches to ensure that changes in projected Lyme disease case counts resulted from predicted changes in incidence rather than projected population growth or decline. We report point estimates and 95% prediction intervals when discussing projected changes in Lyme disease case counts.

### Model validation

To evaluate predictive model accuracy, we compared hindcasted Lyme disease incidence under both emissions scenarios to observed values for 2008 – 2017 (Judge et al. 1985, Clark et al. 2001). We compared model accuracy under varying model specifications to check the robustness of the climate-disease relationships. In the first specification, each regional model contained the predictors (climate, land cover, and non-ecological variables) determined through variable selection (see Methods: Statistical analysis) as well as county and year dummy variables. In the second specification, each regional model contained the same predictors as in the first specification, but only linear versions of the climate predictors were included. This is to assess the sensitivity of our results to the functional form of climate-disease relationships. Under the third specification, regional models contained the same climate and non-climate predictors as in the first specification but no dummy variables. Under the fourth specification, regional models contained all possible climate and non-climate variables, and the county and year dummy variables. Using each of these specifications, we created models of Lyme disease incidence on a training dataset containing a randomly selected 75% subset of counties and years and used the withheld 25% of observations for validation (Hijmans 2012, Caldwell et al. 2016). To evaluate the performance of each model specification, we calculated the root-mean-square error (RMSE) and correlation coefficient between projected and observed Lyme disease incidence for a given county and year between 2008 – 2017 (the years with complete data for all predictors) for each regional model. We also compared estimated average annual incidence to observed average annual incidence for each model specification and each region. We used the modeled climate and land cover data when hindcasting as these datasets were used for Lyme disease projections.

## Results

### Climate and Lyme disease incidence

At least one climate variable was included in the best fit model of Lyme disease incidence for all US regions with vector species present (Table 2). However, the specific climate variable(s) included in the model varied between regions and were often not significant predictors of incidence. As hypothesized, cumulative temperature was a significant, positive predictor in the Northeast, while the number of hot, dry days in May - July was a significant, negative predictor in this region (Table 2). Hot, dry days was also a significant, negative predictor in the Midwest. In the Southeast, daily temperature variability was a significant, positive predictor of incidence. In all other regions, the temperature and/or precipitation variables included in the best fit models were not statistically significant predictors. Further, for all regions, the climate predictors explained relatively little of the variation in Lyme disease incidence compared to the county dummy variables (Table 2). In many cases, quadratic versions of climate predictors were included in the best fit model for a particular region, indicating nonlinearity in climate-disease relationships (Table 2). For example, the number of hot, dry days, total annual precipitation, and temperature variability were all nonlinear predictors in the best fit model for the Northeast.

**Table 2.** Effect of climate and non-climate variables on Lyme disease incidence by region. Only variables included in the best fit model, as determined by variable selection, are shown. The scaled coefficient estimates (Coef.) shown here reflect the standard deviation change in Lyme disease incidence for a one standard deviation change in the climate variable. The coefficients are scaled so that the effects of different variables are directly comparable. The standard errors (SE) shown are clustered by the agricultural statistics district (see Methods: Statistical analysis). Statistically significant (p < 0.05) coefficients are denoted with *.

|  | Northeast |  | Midwest |  | Pacific |  | Pacific Southwest |  | Southwest |  | Southeast |  |
| --- | --- | --- | --- | --- | --- | --- | --- | --- | --- | --- | --- | --- |
| Variable | Coef. | SE | Coef. | SE | Coef. | SE | Coef. | SE | Coef. | SE | Coef. | SE |
| Avg. winter temp. |  |  | -0.073 | 0.237 | -0.967 | 1.039 | 0.119 | 0.172 |  |  |  |  |
| Avg. winter temp.<sup>2</sup> |  |  | 0.381 | 0.253 | 1.268 | 0.894 | 0.391 | 0.403 |  |  |  |  |
| Avg. spring precip. | 0.067 | 0.129 | -0.051 | 0.041 |  |  | 0.089 | 0.089 | -0.998 | 0.836 |  |  |
| Avg. spring precip.<sup>2</sup> | -0.094 | 0.083 |  |  |  |  |  |  |  |  |  |  |
| Hot, dry days | -0.302* | 0.128 | -0.264* | 0.099 |  |  |  |  | 0.151 | 0.137 | -0.029 | 0.022 |
| Hot, dry days <sup>2</sup> | 0.106 | 0.062 | 0.121* | 0.055 |  |  |  |  |  |  |  |  |
| Cumulative temp. | 1.034* | 0.468 |  |  |  |  |  |  | 1.589 | 1.429 | 1.928 | 1.657 |
| Cumulative temp.<sup>2</sup> |  |  |  |  |  |  |  |  | -2.127 | 1.620 | -2.405 | 1.811 |
| Total annual precip. | -0.141 | 0.283 | -0.046 | 0.176 |  |  |  |  | 1.192 | 0.981 |  |  |
| Total annual precip.<sup>2</sup> | 0.183 | 0.229 | -0.010 | 0.115 |  |  |  |  |  |  |  |  |
| Temp. variability | 0.365 | 0.596 |  |  |  |  | 0.112 | 0.954 |  |  | 0.813* | 0.310 |
| Temp. variability <sup>2</sup> | 0.131 | 0.483 |  |  |  |  | 0.224 | 0.488 |  |  | -0.473* | 0.241 |
| Precip. variability |  |  | 0.040 | 0.048 |  |  |  |  | -0.220 | 0.176 |  |  |
| Precip. variability <sup>2</sup> |  |  | 0.012 | 0.019 |  |  |  |  |  |  |  |  |
| Lag 'ticks' search | 0.168* | 0.075 | 0.016 | 0.017 | 0.014 | 0.036 | 0.049 | 0.059 | 0.020 | 0.069 | -0.016 | 0.019 |
| Poverty | -0.055 | 0.087 | 0.046 | 0.072 |  |  |  |  | 0.210 | 0.133 |  |  |
| Percent insured |  |  |  |  |  |  |  |  |  |  | -0.009 | 0.039 |
| Forest cover | 1.988 | 1.283 | -3.966 | 3.896 | -1.515* | 0.763 | -0.365 | 0.513 |  |  | 0.663 | 0.383 |
| Mixed dev. cover |  |  |  |  |  |  |  |  | 1.447 | 1.650 | 1.441* | 0.686 |
| R<sup>2</sup> | 0.728 |  | 0.829 |  | 0.405 |  | 0.327 |  | 0.309 |  | 0.330 |  |
| Model with only climate and dummy variables |  |  |  |  |  |  |  |  |  |  |  |  |
| R<sup>2</sup> | 0.681 |  | 0.768 |  | 0.230 |  | 0.137 |  | 0.112 |  | 0.146 |  |
| Model with only non-climate and dummy variables |  |  |  |  |  |  |  |  |  |  |  |  |
| R<sup>2</sup> | 0.712 |  | 0.820 |  | 0.400 |  | 0.308 |  | 0.258 |  | 0.320 |  |
| Model with only county dummy variable |  |  |  |  |  |  |  |  |  |  |  |  |
| R<sup>2</sup> | 0.606 |  | 0.700 |  | 0.156 |  | 0.114 |  | 0.090 |  | 0.149 |  |
| Model with only year dummy variable |  |  |  |  |  |  |  |  |  |  |  |  |
| R<sup>2</sup> | 0.045 |  | 0.018 |  | 0.028 |  | 0.014 |  | 0.007 |  | 0.010 |  |

### Non-climate predictors and Lyme disease incidence

For all regions, the best fit model of Lyme disease incidence included the 1-year lagged tick search frequency as well one health-seeking predictor and/or a land cover variable (Table 2). Lagged tick search frequency was a significant, positive predictor in the Northeast, and had regionally variable, and non-significant effects in other regions. Poverty was negatively associated with Lyme disease incidence in the Northeast, and positively associated with incidence in the Midwest and Southwest, but was not a significant predictor in any of these models. Health insurance coverage was a non-significant, negative predictor of Lyme disease in the Southeast. Forest cover was included in all regional models except the Southwest, but had regionally variable effects and was only a significant predictor in the Pacific. Mixed development cover was a positive predictor in the Southeast and Southwest, but only significant in the Southeast. The above non-climate predictors were included in each regional model of incidence along with county and year dummy variables. The majority of the variation in incidence for each region was explained by the county dummy variable (Table 2), indicating that there was a great deal of unobserved county-level heterogeneity driving Lyme disease incidence that was captured by the dummy variables. However, the estimated effect sizes of the predictors are the marginal effects of deviations from county- and year-means, meaning the total effect of a given variable, such as forest cover, may be larger if much of the variation is captured by the county fixed effects.

### Model Validation

Under the main model specification, hindcasted Lyme disease incidence matched the observed values with reasonable accuracy in the high incidence regions (Table 3 and Figure S1). In the Northeast and Midwest, the correlations between estimated Lyme disease incidence for a given county and year and the observed incidence were 0.85 and 0.90, respectively. Model accuracy was lower in the Pacific, Pacific Southwest, Southwest, and Southeast, where incidence is much lower (r = 0.40, 0.26, 0.07, 0.32, respectively). However, the estimated annual average Lyme disease incidence (i.e., average incidence for a given region between 2008 – 2017) closely matched the observed annual average for all regions (Table 3). For each region, the estimated incidence was within 13% of the observed incidence, and was within 5% for the Northeast specifically.

**Table 3.** Model validation metrics for four specifications of models of Lyme disease incidence (see Methods: Model validation). The model validation metrics shown are the root-mean-square error (RMSE) and correlation coefficient (r) for estimated versus observed Lyme disease incidence in the testing data sets. The observed and estimated average (± 1 standard deviation) annual Lyme disease incidence is also shown for each region and each model specification. Model validation was performed using data from 2008 – 2017 (the years with complete data for all predictors).

|  |  | Main Model |  |  | Model Spec. 2 |  |  | Model Spec. 3 |  |  | Model Spec. 4 |  |  |
| --- | --- | --- | --- | --- | --- | --- | --- | --- | --- | --- | --- | --- | --- |
|  | Observed annual incidence | Est. annual inc. | RMSE | r | Est. annual inc. | RMSE | r | Est. annual inc. | RMSE | r | Est. annual inc. | RMSE | r |
| NE | 48.9 ± 17.4 | 51.3 ± 13.3 | 38.970 | 0.853 | 51.8 ± 15.6 | 39.138 | 0.851 | 49.4 ± 9.4 | 65.419 | 0.458 | 51.2 ± 13.2 | 38.343 | 0.858 |
| MW | 14.5 ± 3.2 | 12.7 ± 2.1 | 15.709 | 0.903 | 12.6 ± 3.1 | 15.706 | 0.902 | 14.2 ± 4.0 | 29.023 | 0.602 | 12.7 ± 2.1 | 15.49 | 0.906 |
| PC | 0.8 ± 0.3 | 0.8 ± 0.1 | 1.739 | 0.402 | 0.9 ± 0.3 | 1.739 | 0.404 | 0.9 ± 0.1 | 1.777 | 0.282 | 0.8 ± 0.1 | 1.736 | 0.423 |
| PS | 0.9 ± 0.6 | 0.8 ± 0.4 | 1.682 | 0.264 | 0.8 ± 0.4 | 1.682 | 0.268 | 0.8 ± 0.2 | 1.316 | 0.321 | 0.8 ± 0.4 | 1.747 | 0.262 |
| SW | 0.4 ± 0.3 | 0.4 ± 0.2 | 5.169 | 0.071 | 0.4 ± 0.3 | 5.170 | 0.070 | 0.3 ± 0.2 | 5.131 | 0.040 | 0.4 ± 0.2 | 5.157 | 0.086 |
| SE | 0.5 ± 0.2 | 0.5 ± 0.2 | 1.685 | 0.323 | 0.5 ± 0.2 | 1.694 | 0.313 | 0.5 ± 0.2 | 1.725 | 0.172 | 0.5 ± 0.2 | 1.682 | 0.326 |

Model accuracy also varied across the four model specifications (Table 3). In particular, model specifications with dummy variables outperformed (i.e., lower RMSE, higher correlation coefficients) those without. Models including only linear versions of climate predictors (i.e., model specification two) along with non-climate and dummy variables performed similarly to the main model specification but with slightly lower correlation coefficients and higher RMSE in the Northeast and Midwest, where the majority of cases occur. Coefficient estimates and Lyme disease projections using this model specification are shown in Tables S4 and S5. Models including all potential climate and non-climate predictors along with dummy variables had similar accuracy to the main model specification and model specification two (Table 3). The simpler, variable selection-based model specification using nonlinear climate predictors where selected was thus used for the remaining analysis to minimize overfitting and decrease transferability concerns (Allen and Fildes 2001, Wenger et al. 2011, Wenger and Olden 2012), and to achieve the greatest accuracy in high Lyme disease incidence regions.

### Projected Lyme disease incidence

Under the upper climate change scenario (RCP8.5), the number of Lyme disease cases in the Northeast is projected to increase by 23,619 ± 21,607 by 2040 – 2050 and 61,776 ± 27,578 by 2090 – 2100 (Figures 2 and 3, Table 4). Non-significant decreases in the Midwest and increases in the Southeast were also projected under this scenario, and minimal, non-significant changes were projected for other regions (Table 4). By contrast, under the moderate climate change scenario (RCP4.5), no regions were projected to significantly increase or decrease. Non-significant increases in the Midwest, and non-significant increases or decreases, depending on the decade, were projected for the Northeast, with minimal changes elsewhere. Given the regionally variable projections and the large prediction intervals around all point estimates, total US Lyme disease incidence is not projected to change significantly under either climate scenario by 2040 – 2050 or 2090 – 2100 (Table 4). These results indicate that future changes in US Lyme disease burden are highly uncertain, vary strongly by region, and will depend on the degree of future climate change.

**Figure 2.**
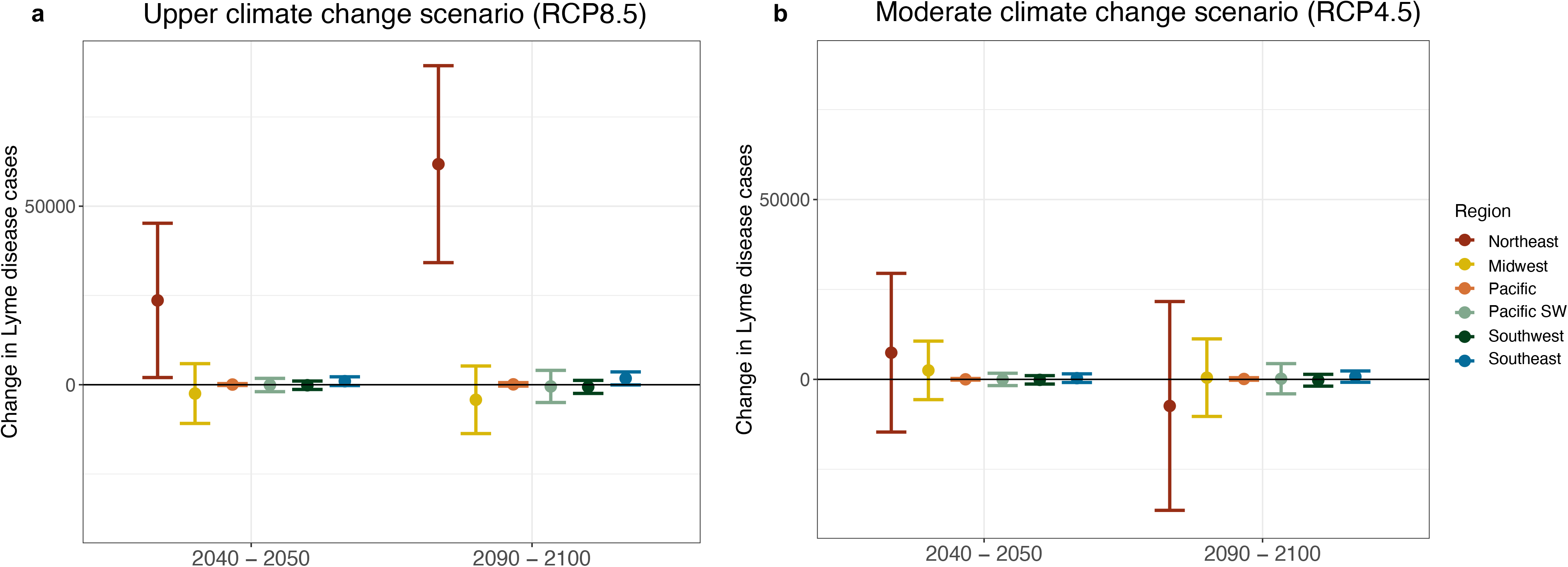
Projected change in Lyme disease cases by region for 2040 – 2050 and 2090 – 2100 under the **a)** upper (RCP8.5) and **b)** moderate (RCP4.5) climate change scenarios. Case changes refer to raw case counts rather than incidence and indicate the average change in cases for a particular decade relative to hindcasted values for 2010 – 2020. Bars represent 95% prediction intervals. Regions are defined in Fig. 1.

**Figure 3.**
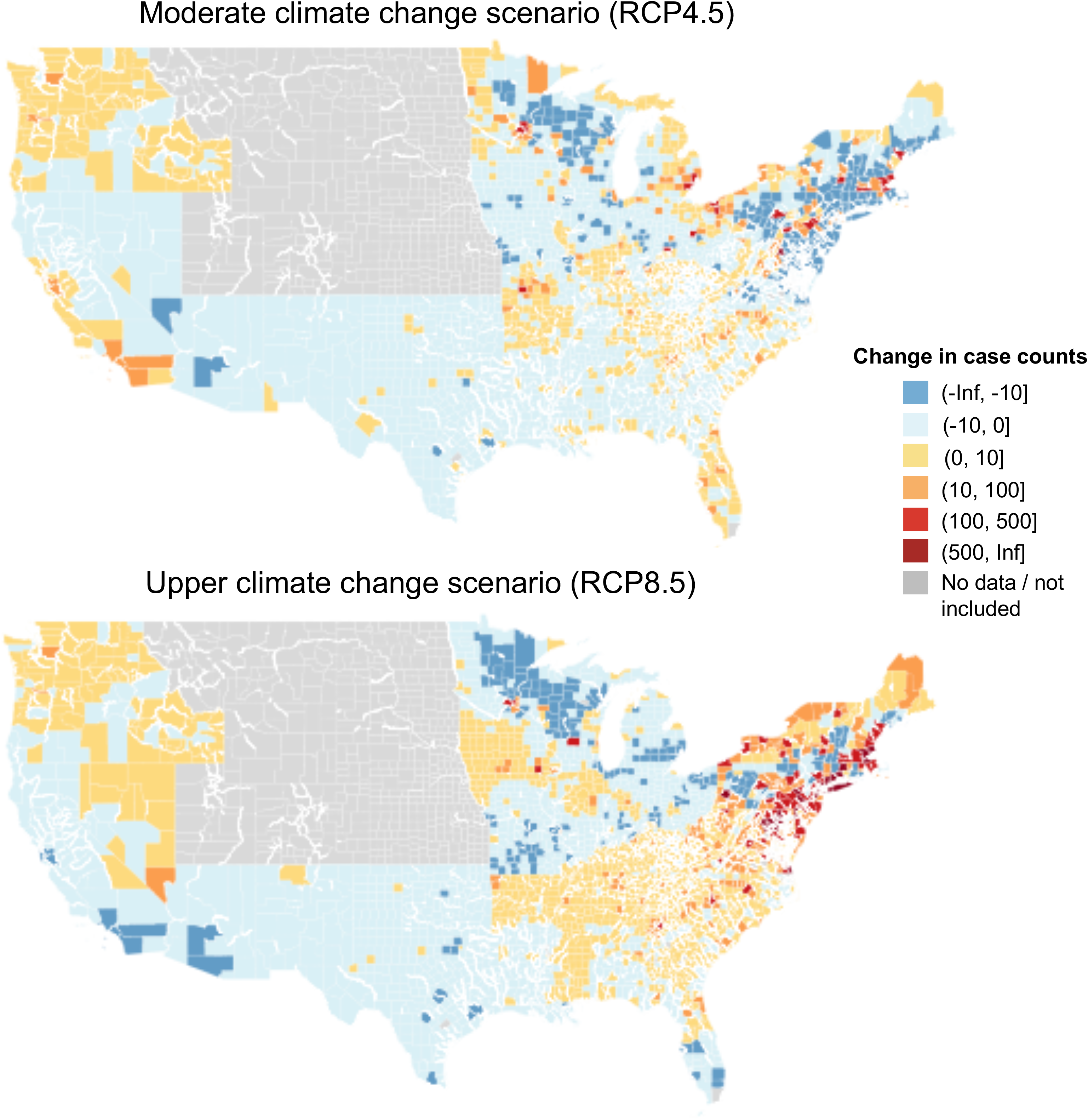
Projected change in Lyme disease cases for 2100 shown at the county level under the **a)** upper (RCP8.5) and **b)** moderate (RCP4.5) climate change scenarios. Case changes refer to raw case counts rather than incidence and are relative to hindcasted values for 2010 – 2020. All counties within the Mountain Prairie are shown in gray as this region was not included in the analysis. Other counties shown in gray (n = 49) containing missing disease, land cover or climate data.

**Table 4.** Projected change in the number of Lyme disease cases, relative to hindcasted 2010 - 2020 levels, for each region under the upper and moderate climate change scenario. Lyme disease projections incorporate county-level population size projections under SSP2 for 2050 and 2100 from Hauer et al. 2019 (see Tables S6 & S7). Point estimates and 95% prediction intervals are shown.

|  | Upper climate change scenario<br>(RCP8.5) |  | Moderate climate change scenario<br>(RCP4.5) |  |
| --- | --- | --- | --- | --- |
|  | 2040 – 2050 | 2090 – 2100 | 2040 – 2050 | 2090 - 2100 |
| Northeast | 23,619<br>[2,013, 45,226] | 61,776<br>[34,197, 89,354] | 7,415<br>[-14,646, 29,476] | -7,385<br>[-36,417, 21,647] |
| Midwest | -2,470<br>[-10,839, 5,899] | -4,217<br>[-13,681, 5,247] | 2,504<br>[-5,633, 10,641] | 477<br>[-10,305, 11,529] |
| Pacific | 48<br>[-218, 315] | 104<br>[-379, 587] | 17<br>[-212, 246] | 113<br>[-246, 471] |
| Pacific Southwest | -84<br>[1,948, 1,780] | -239<br>[-2,490, 2,012] | -11<br>[-1,726, 1,705] | 90<br>[-2,012, 2,192] |
| Southwest | -148<br>[-1325, 1,029] | -608<br>[-2,434, 1,217] | -133<br>[-1,301, 1,034] | -240<br>[-1,884, 1,403] |
| Southeast | 991<br>[-236, 2,217] | 1,768<br>[-61, 3,597] | 339<br>[-865, 1,543] | 776<br>[-807, 2,339] |
| US Total | 22,485<br>[-8,585, 57,451] | 33,639<br>[-9,916, 77,194] | 10,131<br>[-24,383, 44,645] | -6,169<br>[-51,671, 39,581] |

These Lyme disease projections were qualitatively similar to those generated using only linear versions of the climate variables (Table S5). Under this model specification (model specification two, see Methods: Model validation), the number of Lyme disease cases in the Northeast is projected to increase under the upper climate change scenario (21,467 ± 21,354 by 2040 – 2050 and 42,538 ± 24,129 by 2090 – 2100), but not under the moderate climate scenario. Non-significant decreases and increases in the Midwest were projected for the upper and moderate climate scenario, respectively, and non-significant changes in the US as a whole were projected under both scenarios and time periods. These results are all consistent with those generated under the main model specification, indicating that our projections are generally robust to the functional form of climate-disease relationships specified in the model. The one qualitative difference in results is the significant increase in cases in the Southeast under the upper climate change scenario (1,522 ± 1,213 by 2040 – 2050 and 3,460 ± 1,736 by 2090 – 2100) under model specification two, which was marginally non-significant under the main model specification.

Lyme disease case projections made using county-level population size projections were similar to those using constant (i.e., 2017) population sizes. In particular, large but uncertain increases in Lyme diseases cases were still projected for the Northeast under the upper climate change scenario (18,885 ± 19,509 by 2040 – 2050 and 40,320 ± 21,886 by 2090 – 2100) when assuming constant population sizes. This indicates that our results are generally robust to population size assumptions and are not solely driven by projected changes in human demography. However, because population growth is projected for the Northeast (Hauer et al. 2019; Table S7), projections made assuming constant population sizes are smaller (but not significantly) than those using projected population sizes.

## Discussion

Given the increasing rate of vector-borne disease emergence and re-emergence in recent decades, including Zika in Central and South America and tick-borne encephalitis in Europe, identifying the environmental drivers of vector-borne disease transmission has been a major research theme (Rogers and Randolph 2006, Kilpatrick and Randolph 2012, Lafferty and Mordecai 2016, Swei et al. 2019). Extensive prior research indicates that temperature and moisture conditions can impact vector life cycles, activity patterns, abundance, and range limits (reviewed in Ogden and Lindsay 2016). Yet despite clear relationships between specific features of climate and aspects of vector life cycles and biology, identifying how these relationships translate to affect disease incidence has remained challenging. Here we use 18 years of disease and climate data in a panel data statistical modeling approach to identify the impacts of climate change on human Lyme disease incidence across biogeographically distinct US regions. We find that climate was a predictor of interannual variation in Lyme disease incidence in all US regions with established vector species (Northeast, Midwest, Pacific, Pacific Southwest, Southwest, and Southeast), even after controlling for potentially confounding factors and spurious relationships spatially and temporally. However, the specific climate variable(s) that best predicted burdens varied between regions and had highly variable effect sizes and often nonlinear relationships with incidence. While these results underscore the complexity of climate-Lyme disease relationships, the specific associations observed here tended to reflect known relationships between climate and the life histories of the US vectors of Lyme disease, *I. scapularis* and *I. pacificus*.

The strongest climate-disease association detected was between warming annual temperatures and increasing Lyme disease incidence in the Northeast. Previous studies have found that warming year-round temperatures at high latitudes contribute to more rapid tick development rates, increased survival, and *I. scapularis* range expansion (Clow et al. 2017a, Leighton et al. 2012, Lindsay et al. 1995, Ogden et al. 2004, Rand et al. 2004). This suggests warmer temperatures near the ticks’ northern range limit would promote Lyme disease transmission – an expectation empirically supported in this study. We also found a significant negative association between hot, dry conditions during the nymphal questing period (May – July) and incidence in the Northeast and Midwest. Prior studies indicate that desiccating conditions reduce tick questing activity, which can lead to decreased contact rates with larger vertebrate hosts, including humans (Randolph and Storey 1999, Prusinski et al. 2006, Sonenshine and Roe 2013). Further, Burtis et al. 2016 found the number of hot, dry days during this period was significantly negatively associated with *I. scapularis* questing density as well as Lyme disease incidence in the Hudson Valley, Southern New England, and northern New Jersey. Our work thus provides evidence that these prior relationships between desiccating conditions and tick questing behavior scale to incidence across the Northeast and Midwest. That this relationship was not observed or significant in the Southeast or Southwest is also consistent with prior evidence of differing questing behavior in northern and southern *I. scapularis* nymphs. Northern *I. scapularis* nymphs are much more likely to quest above the leaf litter, while southern *I. scapularis* nymphs primarily use habitats below the vegetative surface (Arsnoe et al. 2015). As this different questing behavior buffers southern *I. scapularis* from desiccating conditions, variation in the number of hot, dry days is less likely to impact tick-host contact rates and disease transmission here. Similar differences in questing behavior have been demonstrated between northern and southern population of *I. pacificus* (Lane et al. 2013, MacDonald and Briggs 2016), but we find no significant relationship between hot, dry days and incidence in the Pacific, potentially because low Lyme disease incidence in this region reduces the power to detect effects of variation in climate on incidence. Although we did find the expected negative relationship between hot, dry days and incidence in the Northeast and Midwest, we did not detect the hypothesized positive relationship between spring precipitation and Lyme disease incidence in any region. We did find a positive association in the Northeast and Pacific Southwest, but the association was not significant, and it was negative (but non-significant) in the Midwest and Southwest. This may be due to counteracting effects of precipitation on human behavior leading to reduced tick-human contact rates (Jaenson et al. 2012), independent of effects of precipitation on tick host-seeking suitability.

The associations between climate conditions and Lyme disease incidence found here were detected while rigorously controlling for non-climate predictors of disease as well as unobserved predictors that covary with climate at the county and year levels. In particular, we explicitly controlled for variation in human awareness of ticks, land use and land cover characteristics, proxies for health-seeking behavior, and other unobserved heterogeneity between US counties and years in our modeling approach. Increasing tick awareness, as determined by the frequency of tick-related Google searches, was generally positively associated with Lyme disease incidence, while land cover and health-seeking behavior predictors had regionally variable relationships. By controlling for these effects, we provide strong evidence that the positive association between warming temperatures and Lyme disease incidence in the Northeast found in this study is not simply driven by increasing human awareness of tick-borne disease, temporal trends, or other concurrent changes as has been previously suggested (Morshed et al. 2006, Randolph 2010, Scott and Scott 2018). Further, the total effects of climate and land use predictors may be larger than those estimated here, because these ecological predictors may underlie some of the variation included in the county and year dummy variables.

While our statistical models included both climate and non-climate predictors of Lyme disease incidence, model accuracy varied widely between regions. Most notably, model accuracy was substantially greater for endemic regions (Northeast and Midwest), compared to low incidence (non-endemic) regions (Pacific, Pacific Southwest, Southwest, and Southeast) (Ciesielski et al. 1988). The relatively poor predictive accuracy in non-endemic regions could be due to higher misdiagnosis rates and/or higher travel-associated Lyme disease transmission (Eldin and Parola 2018, Parola and Paddock 2018) decoupling the relationship between local conditions and disease. However, evidence suggests that most Lyme disease transmission occurs in the peri-domestic environment, in which the county of transmission and reporting are likely to be the same (Falco and Fish 1988, Maupin et al. 1991, Jackson et al. 2006, Connally et al. 2009). The lower predictive accuracy in these regions more likely reflects a lack of sufficient annual variation in Lyme disease incidence needed to detect effects of climate in these regions above and beyond the county and year fixed effects, and/or weaker effects of climate conditions on Lyme disease transmission relative to confounding drivers not included in our model such as host movement and community composition. In contrast, the largest effect of climate on disease transmission is expected at the edges of the climate suitability for transmission (Githeko et al. 2000). As portions of the Northeast and Midwest are near the *I. scapularis* northern range limit, the higher model accuracy here likely indicates stronger climate – Lyme disease relationships. Supporting this assertion, the climate predictors explained a relatively larger proportion of the variation in incidence in these regions.

Our Lyme disease projections, made using regionally-specific incidence models and projected climate and land cover data, suggest that climate change may lead to substantial increases in incidence in coming decades, but that these increases are largely concentrated in the Northeast, are highly uncertain, and depend upon the magnitude of climate change. In particular, under the upper climate change scenario (RCP8.5), Lyme disease cases in the Northeast are projected to increase by 23,619 ± 21,607 by 2040 – 2050 and 61,776 ± 27,578 by 2090 – 2100 (Table 4). However, increases are not projected in the Northeast under the moderate climate change scenario (RCP4.5), nor for any other region under either scenario. Large increases in the Midwest under less severe warming are possible, as are large increases in total US cases under more severe warming, but these projections are non-significant. While the significant increase in Lyme disease cases projected for the Northeast under RCP8.5 was robust to alternative model specifications and assumptions about county-level population growth, the large prediction intervals around our point estimates for this region and all others indicate a wide range of potential disease outcomes under climate change.

These results indicate that climate change will likely contribute to increasing Lyme disease incidence in the Northeast, but the specific numerical projections should be interpreted with caution. While significant increases were projected in the Northeast, many other factors contribute to Lyme disease transmission including host movement and community composition, and human avoidance behaviors (Ostfeld 1997, Brownstein et al. 2005b, Ogden et al. 2008, Brinkerhoff et al. 2011, Larsen et al. 2014, Berry et al. 2018, MacDonald et al. 2019a). Accordingly, we found that unobserved county-level heterogeneity, which would encompass these factors, was a predominant driver of incidence in each of our regional models. Thus, while climate may contribute to increasing Lyme disease incidence in northern regions, it may not be the dominant driver of future changes in Lyme disease. Further, while we examined the effects of two potential climate scenarios, uncertainty in these climate change projections was not incorporated into our predictive models and would contribute additional uncertainty in Lyme disease projections. Lastly, the projection models extrapolate from climate and disease relationships observed in the previous 18 years, assuming that these relationships can be extended to climate conditions not yet experienced. That is, we assume that the relationship between cumulative temperature, for example, and Lyme disease incidence in a given region will remain the same even as cumulative temperatures exceed prior values. This could generate inaccurate projections for regions near current tick upper thermal limits such as the Southeast and Southwest as further warming and drought here may reduce tick survival and host-seeking suitability (Vail and Smith 1998, Randolph and Storey 1999, Schulze et al. 2001, Berger et al. 2014, MacDonald et al. 2020). Generating more accurate projections for these regions would require experiments investigating effects of future temperatures on aspects of tick-borne disease transmission.

Despite these limitations and the large uncertainty in our Lyme disease projections, our results are consistent with a growing body of evidence linking increased Lyme disease risk with climate warming (Brownstein et al. 2005a, Burtis et al. 2016, Clow et al. 2017b, Dumic and Severnini 2018, Kilpatrick et al. 2017, Leighton et al. 2012, Ogden et al. 2008,2014b, Robinson et al. 2015, Subak 2003, Tuite et al. 2013). Specifically, our finding of climate change-induced increases in Lyme disease burden at higher latitudes, is consistent with prior studies projecting or observing increasing *I. scapularis* habitat suitability and range expansion under climate warming (Ogden et al. 2008, 2014a, McPherson et al. 2017). Similar range expansions have also been projected and observed for *Ixodes ricinus*, the European Lyme disease vector, under climate warming (Gray et al. 2009, Jaenson and Lindgren 2011, Lindgren et al. 2000, Porretta et al. 2013). Further, our finding that the projected changes in incidence depend on the degree of future warming is also consistent with prior work. *I. scapularis* range expansion and population growth, and the proportion of Eastern Canadians at risk for Lyme disease, are projected to be higher under upper climate change scenarios than under mitigation scenarios (Leighton et al. 2012, McPherson et al. 2017). These results suggest that vector range expansions and future Lyme disease burdens depend in part on climate policy actions.

More generally, our results are consistent with expectations from vector thermal biology that suggest that warming temperatures generally increase transmission near the cold edge of a vector’s range limit, but may decrease or have variable effects elsewhere (Martens et al. 1995, Ogden and Lindsay 2016, Lafferty and Mordecai 2016, Mordecai et al. 2019). For tick-borne diseases, as for other vector-borne diseases, multiple temperature-sensitive traits combine to influence transmission, including survival, development rates, and host-seeking (Randolph et al. 2002, Ogden et al. 2004, Randolph 2004, Ogden and Lindsay 2016, Ogden 2017). Nonlinear effects of temperature on these traits typically leads to vector-borne disease transmission peaking at intermediate temperatures and declining as temperatures approach lower and upper thermal limits (Mordecai et al. 2019). This suggests that climate warming would most strongly increase transmission near the lower thermal limits, such as in the Northeast, as was observed here. This further suggests the effects of climate warming would differ in magnitude and direction depending on the extent of warming, as seen in the Midwest region where non-significant increases were projected under the moderate climate change scenario while decreases were projected under the upper scenario. The theoretical expectations of nonlinear thermal responses therefore help to explain some of the context-dependent effects of temperature found empirically in this study.

## Conclusions

We demonstrate that interannual variation in Lyme disease incidence is associated with climate in all US regions with established vector species, independent of other drivers of disease risk and excluding potentially spurious relationships with county- and year-specific variation. The specific climate variable(s) associated with incidence and their effect sizes varied by region, but the strongest climate-disease association observed was between warming temperatures and increasing incidence in the Northeast. However, in all regions, climate explained less variation in incidence than unobserved county-specific heterogeneity, highlighting that climate is one of many factors influencing Lyme disease transmission. We project that future climate change could substantially increase Lyme disease burden in the Northeast in coming decades under an upper climate change scenario. Cases in the Northeast were not projected to increase under a moderate climate change scenario, highlighting the potential for climate change mitigation to protect human health by preventing further increases in Lyme disease incidence. However, the projected effects in this region and all others are highly uncertain, indicating a wide range of potential disease outcomes under climate change. Our projections provide an essential first step in determining broad patterns of Lyme disease risk under climate change, but ongoing surveillance efforts and mechanistic studies linking changes in vector ecology under climate change to human disease incidence should be conducted to refine these risk assessments.

## Author Contributions

LIC and EAM conceived of the project. All authors designed the analyses. LIC gathered the data and performed the analyses. LIC drafted the manuscript. AJM and EAM revised the manuscript. All authors read and approved the final manuscript. LIC was funded by the Stanford Graduate Fellowship. AJM was funded by a UC Santa Barbara Faculty Research Grant. EAM was funded by an NSF Ecology and Evolution of Infectious Diseases grant (DEB-1518681), the Terman Award, and the NIH NIGMS Maximizing Investigators’ Research Award (R35GM133439).

## Acknowledgements

We are grateful to the CDC Division of Vector-Borne Diseases for supplying Lyme disease case data, Mohammad Alhamdan from NASA for supplying climate data, and to Iain Caldwell, Jamie Caldwell, Marissa Childs, Johannah Farner, Elizabeth Hadly, Morgan Kain, Devin Kirk, Giulio de Leo, Nicole Nova, and Marta Shocket for providing helpful feedback on the manuscript.

## Data Accessibility

All datasets used in this study are free and publicly available. These datasets can be found here: https://github.com/lcouper/LymeDiseaseClimateChange, along with information about where and when they were originally accessed.

## Notes

https://github.com/lcouper/LymeDiseaseClimateChange

### Competing Interest Statement

The authors have declared no competing interest.

### Summary of Updates

The projected Lyme disease cases including population growth projections are now included in the main text while those assuming no population growth are in the supplementals. Similarly, the model specification including nonlinear climate variables has been moved to the main text while the specification including only linear climate variables is now in the supplementals. Diabetes incidence has now been removed from the list of potential non-ecological predictors of Lyme disease transmission. The discussion has been updated to emphasize that climate variables collectively explain less variation in incidence than unobserved county-level heterogeneity.

## References

Adrion, E. R., J. Aucott, K. W. Lemke, and J. P. Weiner. 2015. Health care costs, utilization and patterns of care following Lyme disease. PLOS ONE 10:e0116767.

Allen, P. G., and R. Fildes. 2001. Econometric Forecasting. Pages 303–362 *in* J. S. Armstrong, editor. Principles of Forecasting. Springer US, Boston, MA.

Armstrong, P. M., L. R. Brunet, A. Spielman, and S. R. Telford III. 2001. Risk of Lyme disease: perceptions of residents of a Lone Star tick-infested community. Bulletin of the World Health Organization 79:916–925.

Arsnoe, I. M., G. J. Hickling, H. S. Ginsberg, R. McElreath, and J. I. Tsao. 2015. Different populations of blacklegged tick nymphs exhibit differences in questing behavior that have implications for human Lyme disease risk. PLOS ONE 10:e0127450.

Bascle, G. 2008. Controlling for endogeneity with instrumental variables in strategic management research. Strategic Organization 6:285–327.

Berger, K. A., H. S. Ginsberg, K. D. Dugas, L. H. Hamel, and T. N. Mather. 2014. Adverse moisture events predict seasonal abundance of Lyme disease vector ticks (Ixodes scapularis). Parasites & Vectors 7:181.

Berry, K., J. Bayham, S. R. Meyer, and E. P. Fenichel. 2018. The allocation of time and risk of Lyme: A case of ecosystem service income and substitution effects. Environmental and Resource Economics 70:631–650.

Brinkerhoff, R. J., C. M. Folsom-O’Keefe, K. Tsao, and M. A. Diuk-Wasser. 2011. Do birds affect Lyme disease risk? Range expansion of the vector-borne pathogen *Borrelia burgdorferi*. Frontiers in Ecology and the Environment 9:103–110.

Brownstein, J. S., T. R. Holford, and D. Fish. 2003. A climate-based model predicts the spatial distribution of the Lyme disease vector Ixodes scapularis in the United States. Environmental Health Perspectives 111:1152–1157.

Brownstein, J. S., T. R. Holford, and D. Fish. 2005a. Effect of climate change on Lyme disease risk in North America. EcoHealth 2:38–46.

Brownstein, J. S., D. K. Skelly, T. R. Holford, and D. Fish. 2005b. Forest fragmentation predicts local scale heterogeneity of Lyme disease risk. Oecologia 146:469–475.

Burtis, J. C., P. Sullivan, T. Levi, K. Oggenfuss, T. J. Fahey, and R. S. Ostfeld. 2016. The impact of temperature and precipitation on blacklegged tick activity and Lyme disease incidence in endemic and emerging regions. Parasites & Vectors 9.

Caldwell, J., S. Heron, C. Eakin, and M. Donahue. 2016. Satellite SST-based coral disease outbreak predictions for the Hawaiian archipelago. Remote Sensing 8:93.

Cameron, A. C., and D. L. Miller. 2015. A practitioner’s guide to cluster-robust inference. Journal of Human Resources 50:317–372.

Clark, J. S., S. R. Carpenter, M. Barber, S. Collins, A. Dobson, J. A. Foley, D. M. Lodge, M. Pascual, R. Pielke, W. Pizer, C. Pringle, W. V. Reid, K. A. Rose, O. Sala, W. H. Schlesinger, D. H. Wall, and D. Wear. 2001. Ecological forecasts: An emerging imperative. Science 293:657–660.

Clow, K. M., P. A. Leighton, N. H. Ogden, L. R. Lindsay, P. Michel, D. L. Pearl, and C. M. Jardine. 2017a. Northward range expansion of *Ixodes scapularis* evident over a short timescale in Ontario, Canada. PLOS ONE 12:e0189393.

Clow, K. M., N. H. Ogden, L. R. Lindsay, P. Michel, D. L. Pearl, and C. M. Jardine. 2017b. The influence of abiotic and biotic factors on the invasion of I*xodes scapularis* in Ontario, Canada. Ticks and Tick-borne Diseases 8:554–563.

Dennis, D. T., T. S. Nekomoto, J. C. Victor, W. S. Paul, and J. Piesman. 1998. Reported Distribution of *Ixodes scapularis* and *Ixodes pacificus* (Acari: Ixodidae) in the United States. Journal of Medical Entomology 35:629–638.

Dister, S. W., and D. Fish. 1997. Landscape characterization of peridomestic risk for Lyme disease using satellite imagery. Americal Journal of Tropical Medicine and Hygiene:6.

Dumic, I., and E. Severnini. 2018. “Ticking bomb”: The impact of climate change on the incidence of Lyme disease. 2018:5719081

Eisen, R. J., L. Eisen, N. H. Ogden, and C. B. Beard. 2016. Linkages of weather and climate with *Ixodes scapularis* and *Ixodes pacificus* (Acari: Ixodidae), enzootic transmission of *Borrelia burgdorferi* and Lyme disease in North America. Journal of Medical Entomology 53:250–261.

Estrada-Peña, A. 2002. Increasing habitat suitability in the United States for the tick that transmits Lyme disease: a remote sensing approach. Environmental Health Perspectives 110:635–640.

Frank, D. H., D. Fish, and F. H. Moy. 1998. Landscape features associated with Lyme disease risk in a suburban residential environment. Landscape Ecology 13:27–36.

Gigon, F. 1985. Biologie d’Ixodes ricinus L. sur le Plateau Suisse – une contribution à l’écologie de ce vecteur. Université de Neuchâtel, Neuchâtel, Switzerland.

Ginsberg, H. S., M. Albert, L. Acevedo, M. C. Dyer, I. M. Arsnoe, J. I. Tsao, T. N. Mather, and R. A. LeBrun. 2017. Environmental factors affecting survival of immature *Ixodes scapularis* and implications for geographical distribution of Lyme disease: The climate/behavior hypothesis. PLOS ONE 12:e0168723.

Glass, G. E., B. S. Schwartz, J. M. Morgan, D. T. Johnson, P. M. Noy, and E. Israel. 1995. Environmental risk factors for Lyme disease identified with geographic information systems. American Journal of Public Health 85:944–948.

González, C., O. Wang, S. E. Strutz, C. González-Salazar, V. Sánchez-Cordero, and S. Sarkar. 2010. Climate change and risk of Leishmaniasis in North America: Predictions from ecological niche models of vector and reservoir species. PLoS Neglected Tropical Diseases 4:e585.

Gray, J. S., H. Dautel, A. Estrada-Peña, O. Kahl, and E. Lindgren. 2009. Effects of climate change on ticks and tick-borne diseases in Europe. Interdisciplinary Perspectives on Infectious Diseases 2009:1–12.

Hahn, M. B., C. S. Jarnevich, A. J. Monaghan, and R. J. Eisen. 2016. Modeling the geographic distribution of *Ixodes scapularis* and *Ixodes pacificus* (Acari: Ixodidae) in the contiguous United States. Journal of Medical Entomology 53:1176–1191.

Hair, J. F., W. C. Black, B. Babin, and R. Anderson, editors. 2014. Multivariate data analysis. Seventh edition. Pearson.

Hauer, M. E. 2019. Population projections for U.S. counties by age, sex, and race controlled to shared socioeconomic pathway. Scientific Data 6.

Hayhoe, K., J. Edmonds, R. E. Kopp, A. N. LeGrande, B. M. Sanderson, M. F. Wehner, and D. J. Wuebbles. 2017. 2017: Climate models, scenarios, and projections. Pages 133–160 *in* D. J. Wuebbles, D. W. Fahey, K. A. Hibbard, D. J. Dokken, B. C. Stewart, and T. K. Maycock, editors. Climate Science Special Report: Fourth National Climate Assessment, Volume I. U.S. Global Change Research Program Washington, DC, USA.

Herrmann, C., and L. Gern. 2013. Survival of *Ixodes ricinus* (Acari: Ixodidae) nymphs under cold conditions is negatively influenced by frequent temperature variations. Ticks and Tick-borne Diseases 4:445–451.

Hii, Y. L., J. Rocklöv, N. Ng, C. S. Tang, F. Y. Pang, and R. Sauerborn. 2009. Climate variability and increase in intensity and magnitude of dengue incidence in Singapore. Global Health Action 2:2036.

Hijmans, R. J. 2012. Cross-validation of species distribution models: removing spatial sorting bias and calibration with a null model. Ecology 93:679–688.

Humphrey, P. T., D. A. Caporale, and D. Brisson. 2010. Uncoordinated phylogeography of *Borrelia burgdorferi* and its tick vector, *Ixodes scapularis*. Evolution 64:2653–2663.

Jaenson, T. G., D. G. Jaenson, L. Eisen, E. Petersson, and E. Lindgren. 2012. Changes in the geographical distribution and abundance of the tick *Ixodes ricinus* during the past 30 years in Sweden. Parasites & Vectors 5:1–15.

Jaenson, T. G. T., and E. Lindgren. 2011. The range of *Ixodes ricinus* and the risk of contracting Lyme borreliosis will increase northwards when the vegetation period becomes longer. Ticks and Tick-borne Diseases 2:44–49.

Johnson, L., A. Aylward, and R. B. Stricker. 2011. Healthcare access and burden of care for patients with Lyme disease: A large United States survey. Health Policy 102:64–71.

Jones, C. J., and U. D. Kitron. 2000. Populations of *Ixodes scapularis* (Acari: Ixodidae) are modulated by drought at a Lyme disease focus in Illinois. Journal of Medical Entomology 37:408–415.

Judge, G., W. Griffiths, R. Carter, H. Lutkepohl, and T. Lee. 1985. The theory and practice of econometrics. Wiley, New York.

Kain, D. E., F. A. H. Sperling, H. V. Daly, and R. S. Lane. 1999. Mitochondrial DNA sequence variation in *Ixodes pacificus* (Acari: Ixodidae). Heredity 83:378–386.

Kain, D. E., F. A. H. Sperling, and R. S. Lane. 1997. Population genetic structure of *Ixodes pacificus* (Acari: Ixodidae) using allozymes. Journal of Medical Entomology 34:441–450.

Killilea, M. E., A. Swei, R. S. Lane, C. J. Briggs, and R. S. Ostfeld. 2008. Spatial Dynamics of Lyme Disease: A Review. EcoHealth 5:167–195.

Kilpatrick, A. M., A. D. M. Dobson, T. Levi, D. J. Salkeld, A. Swei, H. S. Ginsberg, A. Kjemtrup, K. A. Padgett, P. M. Jensen, D. Fish, N. H. Ogden, and M. A. Diuk-Wasser. 2017. Lyme disease ecology in a changing world: consensus, uncertainty and critical gaps for improving control. Philosophical Transactions of the Royal Society B: Biological Sciences 372:20160117.

Kilpatrick, A. M., and S. E. Randolph. 2012. Drivers, dynamics, and control of emerging vector-borne zoonotic diseases. The Lancet 380:1946–1955.

Kurtenbach, K., K. Hanincová, J. I. Tsao, G. Margos, D. Fish, and N. H. Ogden. 2006. Fundamental processes in the evolutionary ecology of Lyme borreliosis. Nature Reviews Microbiology 4:660–669.

Lafferty, K. D., and E. A. Mordecai. 2016. The rise and fall of infectious disease in a warmer world. F1000Research 5.

Lane, R. S., N. Fedorova, J. E. Kleinjan, and M. Maxwell. 2013. Eco-epidemiological factors contributing to the low risk of human exposure to ixodid tick-borne borreliae in southern California, USA. Ticks and Tick-borne Diseases 4:377–385.

Lane, R. S., J. E. Kleinjan, and G. B. Schoeler. 1995. Diel activity of nymphal *Dermacentor occidentalis* and *Ixodes pacificus* (Acari: Ixodidae) in relation to meteorological factors and host activity periods. Journal of Medical Entomology 32:290–299.

Larsen, A. E., A. J. MacDonald, and A. J. Plantinga. 2014. Lyme disease risk influences human settlement in the wildland–urban interface: Evidence from a longitudinal analysis of counties in the northeastern United States. The American Journal of Tropical Medicine and Hygiene 91:747–755.

Larsen, A. E., K. Meng, and B. E. Kendall. 2019. Causal analysis in control-impact ecological studies with observational data. Methods in Ecology and Evolution.

Leighton, P. A., J. K. Koffi, Y. Pelcat, L. R. Lindsay, and N. H. Ogden. 2012a. Predicting the speed of tick invasion: an empirical model of range expansion for the Lyme disease vector *Ixodes scapularis* in Canada. Journal of Applied Ecology 49:457–464.

Lindgren E, Tälleklint L, and Polfeldt T. 2000. Impact of climatic change on the northern latitude limit and population density of the disease-transmitting European tick *Ixodes ricinus*. Environmental Health Perspectives 108:119–123.

Lindsay, L. R., I. K. Barker, G. A. Surgeoner, S. A. McEwen, T. J. Gillespie, and J. T. Robinson. 1995. Survival and development of *Ixodes scapularis* (Acari: Ixodidae) under various climatic conditions in Ontario, Canada. Journal of Medical Entomology 32:143–152.

Loevinsohn, M. E. 1994. Climatic warming and increased malaria incidence in Rwanda. The Lancet 343:714–718.

MacDonald, A. J., and C. J. Briggs. 2016. Truncated seasonal activity patterns of the western blacklegged tick (*Ixodes pacificus*) in central and southern California. Ticks and Tick-borne Diseases 7:234–242.

MacDonald, A. J., A. E. Larsen, and A. J. Plantinga. 2019a. Missing the people for the trees: Identifying coupled natural–human system feedbacks driving the ecology of Lyme disease. Journal of Applied Ecology 56:354–364.

MacDonald, A. J., S. McComb, C. O’Neill, K. A. Padgett, and A. E. Larsen. 2020. Projected climate and land use change alter western blacklegged tick phenology, seasonal host-seeking suitability and human encounter risk in California. Global Change Biology 2020:1–16.

MacDonald, A. J., and E. A. Mordecai. 2019. Amazon deforestation drives malaria transmission, and malaria burden reduces forest clearing. Proceedings of the National Academy of Sciences 116:22212–22218.

MacDonald, A. J., C. O’Neill, M. H. Yoshimizu, K. A. Padgett, and A. E. Larsen. 2019b. Tracking seasonal activity of the western blacklegged tick across California. Journal of Applied Ecology 56:2562–2573.

Martens, W., T. Jetten, J. Rotmans, and L. Niessen. 1995. Climate change and vector-borne diseases: A global modelling perspective. Global Environmental Change 5:195–209.

Mattingly, P. F. 1969. The biology of mosquito-borne disease. London: George Allen and Unwin Ltd.

McCabe, G. J., and J. E. Bunnell. 2004. Precipitation and the occurrence of Lyme disease in the northeastern United States. Vector-Borne and Zoonotic Diseases 4:143–148.

McClure, M., and M. A. Diuk-Wasser. 2019. Climate impacts on blacklegged tick host-seeking behavior. International Journal for Parasitology 49:37–47.

McEnroe, W. D. 1977. Restriction of the species range of *Ixodes scapularis*, Say, in Massachusetts by fall and winter temperature. Acarologia.

McPherson, M., A. García-García, F. J. Cuesta-Valero, H. Beltrami, P. Hansen-Ketchum, D. MacDougall, and N. H. Ogden. 2017. Expansion of the Lyme disease vector *Ixodes scapularis* in Canada inferred from CMIP5 climate projections. Environmental Health Perspectives 125:057008.

Mills, J. N., K. L. Gage, and A. S. Khan. 2010. Potential influence of climate change on vector-borne and zoonotic diseases: A review and proposed research plan. Environmental Health Perspectives 118:1507–1514.

Mordecai, E. A., J. M. Caldwell, M. K. Grossman, C. A. Lippi, L. R. Johnson, M. Neira, J. R. Rohr, S. J. Ryan, V. Savage, M. S. Shocket, R. Sippy, A. M. S. Ibarra, M. B. Thomas, and O. Villena. 2019. Thermal biology of mosquito-borne disease. Ecology Letters 22:1690–1708.

Morshed, M. G., J. D. Scott, K. Fernando, G. Geddes, A. Mcnabb, S. Mak, and L. A. Durden. 2006. Distribution and characterization of *Borrelia burgdorferi* isolates from I*xodes scapularis* and presence in mammalian hosts in Ontario, Canada. Journal of Medical Entomology 43:762–773.

Nakicenovic, N., J. Alcamo, A. Grubler, K. Riahi, R. A. Roehrl, H.-H. Rogner, and N. Victor. 2000. Special Report on Emissions Scenarios (SRES), A Special Report of Working Group III of the Intergovernmental Panel on Climate Change. Cambridge University Press, Cambridge.

Nieto, N. C., E. A. Holmes, and J. E. Foley. 2010. Survival rates of immature *Ixodes pacificus* (Acari: Ixodidae) ticks estimated using field-placed enclosures. Journal of Vector Ecology 35:43–49.

Ogden, N. H. 2017. Climate change and vector-borne diseases of public health significance. FEMS Microbiology Letters 364.

Ogden, N. H., M. Bigras-Poulin, C. J. O’Callaghan, I. K. Barker, L. R. Lindsay, A. Maarouf, K. E. Smoyer-Tomic, D. Waltner-Toews, and D. Charron. 2005. A dynamic population model to investigate effects of climate on geographic range and seasonality of the tick *Ixodes scapularis*. International Journal for Parasitology 35:375–389.

Ogden, N. H., and L. R. Lindsay. 2016. Effects of climate and climate change on vectors and vector-borne diseases: Ticks are different. Trends in Parasitology 32:646–656.

Ogden, N. H., L. R. Lindsay, G. Beauchamp, D. Charron, A. Maarouf, C. J. O’Callaghan, D. Waltner-Toews, and I. K. Barker. 2004. Investigation of relationships between temperature and developmental rates of tick *Ixodes scapularis* (Acari: Ixodidae) in the laboratory and field. Journal of Medical Entomology 41:622–633.

Ogden, N. H., A. Maarouf, I. K. Barker, M. Bigras-Poulin, L. R. Lindsay, M. G. Morshed, C. J. O’Callaghan, F. Ramay, D. Waltner-Toews, and D. F. Charron. 2006. Climate change and the potential for range expansion of the Lyme disease vector *Ixodes scapularis* in Canada. International Journal for Parasitology 36:63–70.

Ogden, N. H., M. Radojevic’, X. Wu, V. R. Duvvuri, P. A. Leighton, and J. Wu. 2014a. Estimated effects of projected climate change on the basic reproductive number of the Lyme disease vector *Ixodes scapularis*. Environmental Health Perspectives 122:631–638.

Ogden, N. H., L. St-Onge, I. K. Barker, S. Brazeau, M. Bigras-Poulin, D. F. Charron, C. M. Francis, A. Heagy, R. Lindsay, A. Maarouf, P. Michel, F. Milord, C. J. O’Callaghan, L. Trudel, and A. Thompson. 2008. Risk maps for range expansion of the Lyme disease vector, *Ixodes scapularis*, in Canada now and with climate change. International Journal of Health Geographics 7:24.

Ogden, N., J. Koffi, Y. Pelcat, and L. Lindsay. 2014b. Environmental risk from Lyme disease in central and eastern Canada: a summary of recent surveillance information. Canada Communicable Disease Report 40:74–82.

Ostfeld, R., and J. Brunner. 2015. Climate change and *Ixodes* tick-borne diseases of humans. Philosophical Transactions of the Royal Society B: Biological Sciences 370:20140051.

Ostfeld, R. S. 1997. The ecology of Lyme-disease risk: Complex interactions between seemingly unconnected phenomena determine risk of exposure to this expanding disease. American Scientist 85:338–346.

Padgett, K. A., and R. S. Lane. 2001. Life cycle of *Ixodes pacificus* (Acari: Ixodidae): Timing of developmental processes under field and laboratory conditions. Journal of Medical Entomology 38:684–693.

Peavey, C. A., and R. S. Lane. 1996. Field and laboratory studies on the timing of oviposition and hatching of the western black-legged tick, *Ixodes pacificus* (Acari: Ixodidae). Experimental and Applied Acarology 20:695–711.

Porretta, D., V. Mastrantonio, S. Amendolia, S. Gaiarsa, S. Epis, C. Genchi, C. Bandi, D. Otranto, and S. Urbanelli. 2013. Effects of global changes on the climatic niche of the tick *Ixodes ricinus* inferred by species distribution modelling. Parasites & Vectors 6:271.

Prusinski, M. A., H. Chen, J. M. Drobnack, S. J. Kogut, R. G. Means, J. J. Howard, J. Oliver, G. Lukacik, P. B. Backenson, and D. J. White. 2006. Habitat structure associated with *Borrelia burgdorferi* prevalence in small mammals in New York state. Environmental Entomology 35:308–319.

Purse, B. V., P. S. Mellor, D. J. Rogers, A. R. Samuel, P. P. C. Mertens, and M. Baylis. 2005. Climate change and the recent emergence of bluetongue in Europe. Nature Reviews Microbiology 3:171–181.

Raghavan, R. K., K. Almes, D. G. Goodin, J. A. Harrington, and P. W. Stackhouse. 2014. Spatially heterogeneous land cover/land use and climatic risk factors of tick-borne feline cytauxzoonosis. Vector-Borne and Zoonotic Diseases 14:486–495.

Rand, P. W., M. S. Holman, C. Lubelczyk, E. H. Lacombe, A. T. DeGaetano, and R. P. Smith. 2004. Thermal accumulation and the early development of *Ixodes scapularis*. Journal of Vector Ecology:13.

Randolph, S. E. 1997. Abiotic and biotic determinants of the seasonal dynamics of the tick *Rhipicephalus appendiculatus* in South Africa. Medical and Veterinary Entomology 11:25–37.

Randolph, S. E. 2004. Tick ecology: processes and patterns behind the epidemiological risk posed by ixodid ticks as vectors. Parasitology 129:S37–S65.

Randolph, S. E. 2010. To what extent has climate change contributed to the recent epidemiology of tick-borne diseases? Veterinary Parasitology 167:92–94.

Randolph, S. E., R. M. Green, A. N. Hoodless, and M. F. Peacey. 2002. An empirical quantitative framework for the seasonal population dynamics of the tick *Ixodes ricinus*. International Journal for Parasitology 32:979–989.

Randolph, S. E., and K. Storey. 1999. Impact of microclimate on immature tick-rodent host interactions (Acari: Ixodidae): Implications for parasite transmission. Journal of Medical Entomology 36:741–748.

Ricketts, T. H., E. Dinerstein, D. M. Olson, W. Eichbaum, C. J. Loucks, K. Kavanaugh, P. Hedao, P. Hurley, D. DellaSalla, R. Abell, K. Carney, and S. Walters. 1999. Terrestrial ecoregions of North America: A Conservation Assessment. Island Press.

Rizzoli, A., H. Hauffe, G. Carpi, G. Vourc’h, M. Neteler, and R. Rosa. 2011. Lyme borreliosis in Europe. Euro Surveillance 16.

Robinson, S. J., D. F. Neitzel, R. A. Moen, M. E. Craft, K. E. Hamilton, L. B. Johnson, D. J. Mulla, U. G. Munderloh, P. T. Redig, K. E. Smith, C. L. Turner, J. K. Umber, and K. M. Pelican. 2015. Disease risk in a dynamic environment: The spread of tick-borne pathogens in Minnesota, USA. EcoHealth 12:152–163.

Rodgers, S. E., C. P. Zolnik, and T. N. Mather. 2007. Duration of exposure to suboptimal atmospheric moisture affects nymphal blacklegged tick survival. Journal of Medical Entomology 44:372–375.

Rogelj, J., M. Meinshausen, and R. Knutti. 2012. Global warming under old and new scenarios using IPCC climate sensitivity range estimates. Nature Climate Change 2:248–253.

Rogers, D. J., and S. E. Randolph. 2006. Climate change and vector-borne diseases. Pages 345–381 Advances in Parasitology. Elsevier.

Roiz, D., M. Neteler, C. Castellani, D. Arnoldi, and A. Rizzoli. 2011. Climatic Factors Driving Invasion of the tiger mosquito (*Aedes albopictus*) into new areas of Trentino, Northern Italy. PLoS ONE 6:e14800.

Rosenberg, R., N. P. Lindsey, M. Fischer, C. J. Gregory, A. F. Hinckley, P. S. Mead, G. Paz-Bailey, S. H. Waterman, N. A. Drexler, G. J. Kersh, H. Hooks, S. K. Partridge, S. N. Visser, C. B. Beard, and L. R. Petersen. 2018. Vital Signs: Trends in Reported Vectorborne Disease Cases — United States and Territories, 2004–2016. Morbidity and Mortality Weekly Report 67:496–501.

Salkeld, D. J., and R. S. Lane. 2010. Community ecology and disease risk: lizards, squirrels, and the Lyme disease spirochete in California, USA. Ecology 91:293–298.

Samir, K. C, and Lutz, W. 2017. The human core of the shared socioeconomic pathways: Population scenarios by age, sex and level of education for all countries to 2100. Global Environmental Change 42:181–192.

Schauber, E. M., R. S. Ostfeld, and A. S. E. Jr. 2005. What is the best predictor of annual Lyme disease incidence: Weather, mice, or acorns? Ecological Applications 15:575–586.

Schmidt, G. A., M. Kelley, L. Nazarenko, R. Ruedy, G. L. Russell, I. Aleinov, M. Bauer, S. E. Bauer, M. K. Bhat, R. Bleck, V. Canuto, Y.-H. Chen, Y. Cheng, T. L. Clune, A. Del Genio, R. de Fainchtein, G. Faluvegi, J. E. Hansen, R. J. Healy, N. Y. Kiang, D. Koch, A. A. Lacis, A. N. LeGrande, J. Lerner, K. K. Lo, E. E. Matthews, S. Menon, R. L. Miller, V. Oinas, A. O. Oloso, J. P. Perlwitz, M. J. Puma, W. M. Putman, D. Rind, A. Romanou, M. Sato, D. T. Shindell, S. Sun, R. A. Syed, N. Tausnev, K. Tsigaridis, N. Unger, A. Voulgarakis, M.-S. Yao, and J. Zhang. 2014. Configuration and assessment of the GISS ModelE2 contributions to the CMIP5 archive. Journal of Advances in Modeling Earth Systems 6:141–184.

Schulze, T. L., R. A. Jordan, and R. W. Hung. 2001. Effects of selected meteorological factors on diurnal questing of *Ixodes scapularis* and *Amblyomma americanum* (Acari: Ixodidae). Journal of Medical Entomology 38:318–324.

Scott, J., and C. Scott. 2018. Lyme disease propelled by *Borrelia burgdorferi-infected* blacklegged ticks, wild birds and public awareness — not climate change. Journal of Veterinary Science & Medicine 6:01–08.

Smith, J. R., A. D. Letten, P.-J. Ke, C. B. Anderson, J. N. Hendershot, M. K. Dhami, G. A. Dlott, T. N. Grainger, M. E. Howard, B. M. L. Morrison, D. Routh, P. A. S. Juan, H. A. Mooney, E. A. Mordecai, T. W. Crowther, and G. C. Daily. 2018. A global test of ecoregions. Nature Ecology & Evolution 2:1889–1896.

Sohl, T. 2019. The next generation of land-cover projections for the conterminous United States.

Sohl, T. L., K. L. Sayler, M. A. Bouchard, R. R. Reker, A. M. Friesz, S. L. Bennett, B. M. Sleeter, R. R. Sleeter, T. Wilson, C. Soulard, M. Knuppe, and T. V. Hofwegen. 2014. Spatially explicit modeling of 1992–2100 land cover and forest stand age for the conterminous United States. Ecological Applications 24:1015–1036.

Sonenshine, D. E., and R. M. Roe. 2013. Biology of Ticks. OUP USA.

Stafford, K. C. 1994. Survival of immature *Ixodes scapularis* (Acari: Ixodidae) at different relative humidities. Journal of Medical Entomology 31:310–314.

Subak, S. 2003. Effects of climate on variability in Lyme disease incidence in the Northeastern United States. American Journal of Epidemiology 157:531–538.

Swei, A., L. I. Couper, L. L. Coffey, D. Kapan, and S. Bennett. 2019. Patterns, drivers, and challenges of vector-borne disease emergence. Vector-Borne and Zoonotic Diseases.

Tabachnick, W. J. 2010. Challenges in predicting climate and environmental effects on vector-borne disease episystems in a changing world. Journal of Experimental Biology 213:946–954.

Taylor, K. E., R. J. Stouffer, and G. A. Meehl. 2012. An overview of CMIP5 and the experiment design. Bulletin of the American Meteorological Society 93:485–498.

Tuite, A. R., A. L. Greer, and D. N. Fisman. 2013. Effect of latitude on the rate of change in incidence of Lyme disease in the United States. CMAJ Open 1:E43–E47.

Vail, S. G., and G. Smith. 1998. Air temperature and relative humidity effects on behavioral activity of blacklegged tick (Acari: Ixodidae) nymphs in New Jersey. Journal of Medical Entomology 35:1025–1028.

Vandyk, J. K., D. M. Bartholomew, W. A. Rowley, and K. B. Platt. 1996. Survival of *Ixodes scapularis* (Acari: Ixodidae) exposed to cold. Journal of Medical Entomology 33:6–10.

van Vuuren, D. P., J. Edmonds, M. Kainuma, K. Riahi, A. Thomson, K. Hibbard, G. C. Hurtt, T. Kram, V. Krey, J.-F. Lamarque, T. Masui, M. Meinshausen, N. Nakicenovic, S. J. Smith, and S. K. Rose. 2011. The representative concentration pathways: an overview. Climatic Change 109:5–31.

Wenger, S. J., D. J. Isaak, J. B. Dunham, K. D. Fausch, C. H. Luce, H. M. Neville, B. E. Rieman, M. K. Young, D. E. Nagel, D. L. Horan, and G. L. Chandler. 2011. Role of climate and invasive species in structuring trout distributions in the interior Columbia River Basin, USA. Canadian Journal of Fisheries and Aquatic Sciences 68:988–1008.

Wenger, S. J., and J. D. Olden. 2012. Assessing transferability of ecological models: an underappreciated aspect of statistical validation. Methods in Ecology and Evolution 3:260–267.

Wilking, H., and K. Stark. 2014. Trends in surveillance data of human Lyme borreliosis from six federal states in eastern Germany, 2009–2012. Ticks and Tick-borne Diseases 5:219–224.

Wimberly, M. C., M. J. Yabsley, A. D. Baer, V. G. Dugan, and W. R. Davidson. 2008. Spatial heterogeneity of climate and land-cover constraints on distributions of tick-borne pathogens. Global Ecology and Biogeography 17:189–202.

World Health Organization. 2014. A global brief on vector-borne diseases. World Health Organization Technical Report.

Yamashita, T., K. Yamashita, and R. Kamimura. 2007. A stepwise AIC method for variable selection in linear regression. Communications in Statistics – Theory and Methods 36:2395–2403.

Yang, L., S. Jin, P. Danielson, C. Homer, L. Gass, S. M. Bender, A. Case, C. Costello, J. Dewitz, J. Fry, M. Funk, B. Granneman, G. C. Liknes, M. Rigge, and G. Xian. 2018. A new generation of the United States National Land Cover Database: Requirements, research priorities, design, and implementation strategies. ISPRS Journal of Photogrammetry and Remote Sensing 146:108–123.

Zhang, Z. 2016. Variable selection with stepwise and best subset approaches. Annals of Translational Medicine 4:136–136.

