## Supporting Info for "Impact of prior and projected climate change on US Lyme disease incidence"

### Supporting Information

#### Methods

##### Data availability

All datasets used in this study are free and publicly available. These datasets are available on Github (https://github.com/lcouper/LymeDiseaseClimateChange) along with information about where and when they were originally accessed. Additional details and/or limitations for datasets not fully described in the main text are provided below.

##### Lyme disease case data

Under-, over-, and mis-diagnoses are typical issues in surveillance data (Marier 1977), and particularly problematic for Lyme disease surveillance (Bakken et al. 1992, Steere et al. 1993, Orloski et al. 2000, Naleway et al. 2002, Ertel et al. 2012, Hinckley et al. 2014, Nelson et al. 2015). Testing accuracy and reporting practices vary spatially as regions differ in their infrastructure for Lyme disease surveillance (Bacon et al. 2008). We addressed these issues by explicitly including measures of health-seeking behavior in the modeling approach (described in Methods: Statistical approach) to account for variation in reporting. We also included 'county' and 'year' as dummy variables in the model to account for any additional unobserved heterogeneity amongst counties or between years.

##### Modeled climate data

We used NASA Goddard Institute for Space Studies global climate model ensemble data, accessed through the Earth System Grid Federation, to obtain climate estimates through 2100. We obtained modeled, daily data on near-surface air temperature and precipitation for two climate change scenarios: RCP8.5 and RCP4.5. These scenarios reflect the increase in radiative forcing at the tropopause by 2100 relative to preindustrial levels (8.5 $W/m^2$ and 4.5 $W/m^2$, respectively) (Riahi et al. 2011, Thomson et al. 2011). RCP8.5 reflects the upper range on the open literature on emissions, resulting in substantial increases in greenhouse gas emissions and concentrations, while RCP4.5 reflects a midrange climate policy scenario (Hayhoe et al. 2017). Climate trajectories are derived by working backwards from the radiative forcing values (Hayhoe et al. 2017), meaning there may be slight variation in hindcasted and current climate conditions estimated by the two scenarios. Modeled climate data are available at a 2° x 2.5° resolution. We applied these modeled climate values to counties using the latitude and longitude of NOAA weather stations within a county. Given the coarser spatial resolution of modeled climate data, an average of 3.85 neighboring counties shared a modeled temperature or precipitation value for a given day.

##### Awareness data

The Google Trends dataset supplies the number of searches for a particular term in a given region and time period divided by the total number of searches in that region and time period then multiplied by 100. We initially used the search terms "ticks", "tick bite", and "Lyme disease” as potential indicators of awareness of tick-borne disease. As these search terms generated nearly identical coefficient estimates, we proceeded to use only the “ticks” search term as a predictor. We used “ticks” rather than “tick” as the singular version yielded unrelated search terms (e.g., “tick fire”, “tick tock”).

**Modeled land cover data**

We used land cover projections generated by the USGS Earth Resources Observation and Science Center (EROS) using the Intergovernmental Panel on Climate Change (IPCC) Special Report on Emissions Scenarios (SRES) (Sohl et al. 2014). Annual land cover projections were available through 2100 at a 250-m pixel spatial resolution. We used the SRES A1B and B1 projections to align with the upper and moderate climate change scenarios used in the climate projections (Nakicenovic et al. 2000, Rogelj et al. 2012). Land cover projections included 17 land use and land cover classes from which we calculated two annual, county-level metrics: percent forest cover and percent mechanically disturbed land. Forest cover included the percent evergreen forest, mixed forest, and deciduous forest land cover classes. These land cover classes were available in both the projected and historical data. Mechanically disturbed land included the percent ‘mechanically disturbed private’ and ‘mechanically disturbed other public lands’ classes. These land cover classes refer to private or public land undergoing human-induced changes such as forest clear-cutting or earthmoving. These land cover classes were selected to align with the ‘mixed development’ land cover class, defined as areas with a mixture of constructed materials and vegetation, which was available in the historical but not the projected land cover data.

#### References


Bacon, R.M., Kiersten, K.J. & Mead, P.S. (2008). Surveillance for Lyme Disease - United States, 1992-2006. *MMWR Morb Mortal Wkly Rep*, 57.

Bakken, L.L., Case, K.L., Callister, S.M., Bourdeau, N.J. & Schell, R.F. (1992). Performance of 45 laboratories participating in a proficiency testing program for Lyme disease serology. *JAMA*, 268, 891–895.

Ertel, S.-H., Nelson, R.S. & Cartter, M.L. (2012). Effect of surveillance method on reported characteristics of Lyme disease, Connecticut, 1996–2007. *Emerg. Infect. Dis.*, 18, 242–247.

Hayhoe, K., Edmonds, J., Kopp, R.E., LeGrande, A.N., Sanderson, B.M., Wehner, M.F., *et al.* (2017). 2017: Climate models, scenarios, and projections. In: *Climate Science Special Report: Fourth National Climate Assessment, Volume I* (eds. Wuebbles, D.J., Fahey, D.W., Hibbard, K.A., Dokken, D.J., Stewart, B.C. & Maycock, T.K.). U.S. Global Change Research Program. Washington, DC, USA, pp. 133–160.

Hinckley, A.F., Connally, N.P., Meek, J.I., Johnson, B.J., Kemperman, M.M., Feldman, K.A., *et al.* (2014). Lyme disease testing by large commercial laboratories in the United States. *Clin. Infect. Dis.*, 59, 676–681.

Marier, R. (1977). The reporting of communicable diseases. *Am. J. Epidemiol.*, 105, 587–590.

Nakicenovic, N., Alcamo, J., Grubler, A., Riahi, K., Roehrl, R.A., Rogner, H.-H., *et al.* (2000). *Special Report on Emissions Scenarios (SRES), A Special Report of Working Group III of the Intergovernmental Panel on Climate Change*. Cambridge University Press, Cambridge.

Naleway, A.L., Belongia, E.A., Kazmierczak, J.J., Greenlee, R.T. & Davis, J.P. (2002). Lyme Disease incidence in Wisconsin: a comparison of state-reported rates and rates from a population-based cohort. *Am. J. Epidemiol.*, 155, 1120–1127.

Nelson, C., Saha, S., Kugeler, K., Delorey, M., Shankar, M., Hinckley, A.F., *et al.* (2015). Incidence of clinician-diagnosed Lyme disease, United States, 2005–2010 - Volume 21, Number 9—September 2015 - Emerging Infectious Diseases journal - CDC. *Emerg. Infect. Dis.*

Orloski, K.A., Hayes, E.B., Campbell, G.L. & Dennis, D.T. (2000). Surveillance for Lyme Disease --- United States, 1992- -1998, 11.

Riahi, K., Rao, S., Krey, V., Cho, C., Chirkov, V., Fischer, G., *et al.* (2011). RCP 8.5—A scenario of comparatively high greenhouse gas emissions. *Clim. Change*, 109, 33.

Rogelj, J., Meinshausen, M. & Knutti, R. (2012). Global warming under old and new scenarios using IPCC climate sensitivity range estimates. *Nat. Clim. Change*, 2, 248–253.

Sohl, T.L., Sayler, K.L., Bouchard, M.A., Reker, R.R., Friesz, A.M., Bennett, S.L., *et al.* (2014). Spatially explicit modeling of 1992–2100 land cover and forest stand age for the conterminous United States. *Ecol. Appl.*, 24, 1015–1036.

Steere, A.C., Taylor, E., McHugh, G.L. & Logigian, E.L. (1993). The overdiagnosis of Lyme disease. *JAMA*, 269, 1812–1816.

Thomson, A.M., Calvin, K.V., Smith, S.J., Kyle, G.P., Volke, A., Patel, P., *et al.* (2011). RCP4.5: a pathway for stabilization of radiative forcing by 2100. *Clim. Change*, 109, 77.

#### Figures and Tables

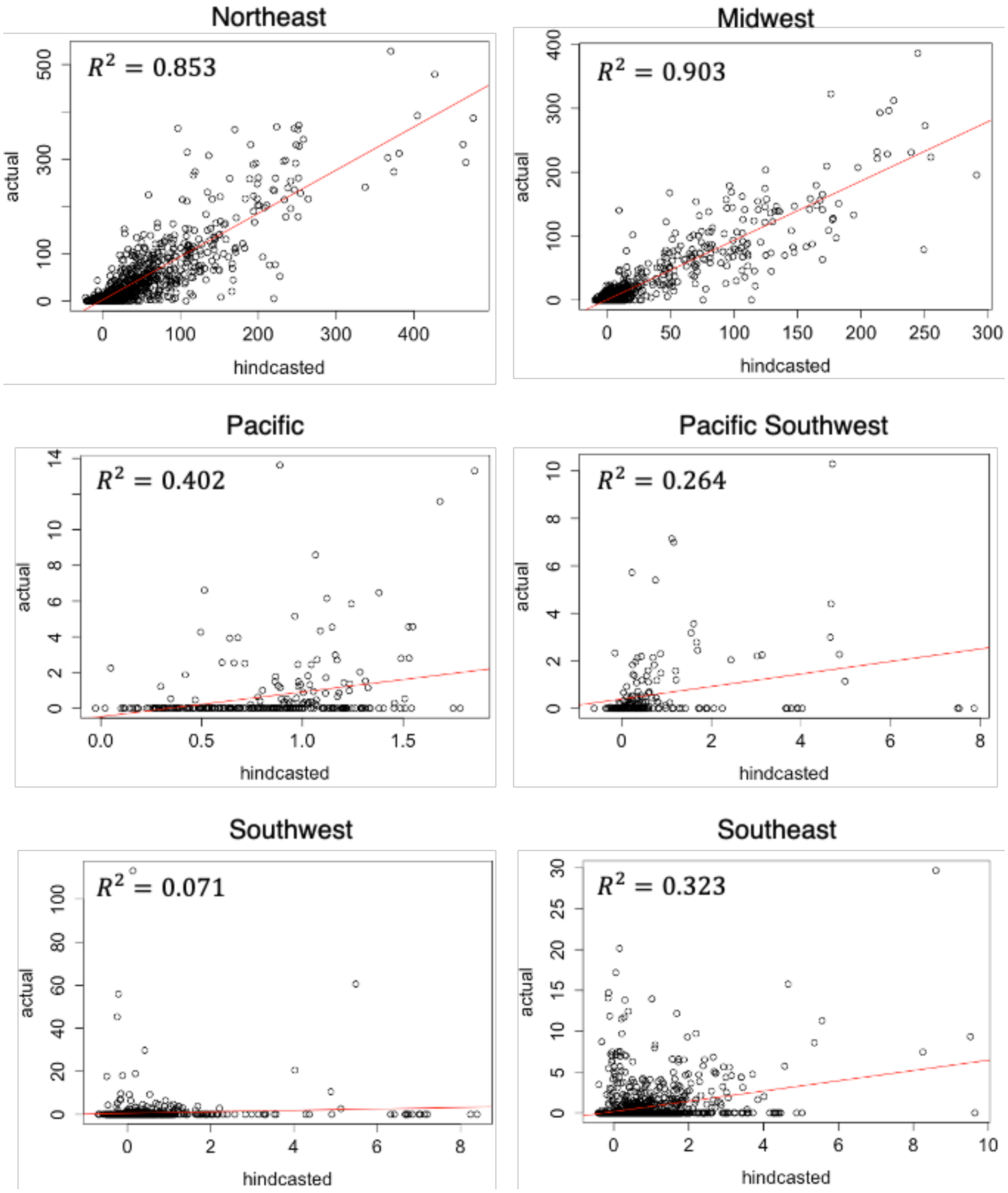


**Figure S1.** Observed and hindcasted Lyme disease incidence by region from 2008 – 2017. The best fit line and correlation coefficient are shown in the plot for each region. Hindcasted Lyme disease incidence was calculated using the main model specification (see Methods: Model validation).

**Table S1.** Hindcasted and projected climate conditions for each region. Only regions included in the analysis are shown. Within each row, values (mean and standard deviation) for climate variables calculated from the upper climate change scenario (RCP8.5) and the moderate climate change scenario (RCP4.5) are listed on top and bottom, respectively.

| | Northeast | Midwest | Pacific | Pacific Southwest | Southwest | Southeast |
|---|---|---|---|---|---|---|
| **Hindcasted Climate (2007 – 2017)** | | | | | | |
| Avg. winter temp. (°F) | 32.4 ± 5.3<br>32.8 ± 5.7 | 28.1 ± 5.9<br>27.6 ± 6.8 | 35.2 ± 8.7<br>34.5 ± 8.8 | 44.1 ± 9.5<br>44.1 ± 9.4 | 47.1 ± 7.0<br>46.8 ± 7.1 | 44.7 ± 7.5<br>44.7 ± 7.5 |
| Avg. spring precip. (mm) | 4.6 ± 1.6<br>4.5 ± 1.6 | 4.8 ± 1.4<br>4.8 ± 1.4 | 3.0 ± 1.3<br>3.2 ± 1.4 | 1.9 ± 1.0<br>2.3 ± 1.2 | 3.1 ± 1.8<br>3.9 ± 1.8 | 4.1 ± 1.5<br>4.4 ± 1.7 |
| Hot, dry days (# days) | 2.7 ± 4.3<br>3.5 ± 4.5 | 3.9 ± 4.5<br>2.9 ± 3.0 | 0.3 ± 0.9<br>0.6 ± 1.4 | 2.6 ± 4.1<br>3.1 ± 4.6 | 24.9 ± 14.8<br>19.0 ± 12.0 | 13.3 ± 10.6<br>12.6 ± 10.1 |
| Cumulative temp (°F) | 19,454 ± 1,457<br>19,535 ± 1,410 | 19,196 ± 1,460<br>19,014 ± 1,441 | 17,440 ± 2,025<br>17,561 ± 2,100 | 20,547 ± 2,499<br>20,463 ± 2,483 | 23,754 ± 2,048<br>23,325 ± 2,092 | 23,145 ± 1,830<br>22,941 ± 1,831 |
| Total annual precip. (mm) | 1,336 ± 299<br>1,303 ± 259 | 1,166 ± 192<br>1,180 ± 228 | 1,432 ± 695<br>1,464 ± 694 | 1,387 ± 565<br>1,469 ± 597 | 863 ± 257<br>958 ± 273 | 1,343 ± 247<br>1,379 ± 270 |
| Temp. var. ($°F^2$) | 269.0 ± 43.2<br>270.1 ± 54.9 | 350.7 ± 60.2<br>350.9 ± 82.7 | 163.3 ± 61.0<br>164.6 ± 62.5 | 142.0 ± 51.4<br>137.8 ± 54.1 | 222.9 ± 56.1<br>208.1 ± 52.1 | 218.1 ± 55.5<br>212.9 ± 58.4 |
| Precip. var. ($mm^2$) | 46.3 ± 16.3<br>45.0 ± 15.1 | 41.8 ± 11.8<br>41.2 ± 13.9 | 35.9 ± 28.0<br>35.3 ± 26.3 | 58.4 ± 37.9<br>64.9 ± 41.8 | 35.9 ± 18.6<br>41.0 ± 21.3 | 57.7 ± 19.6<br>57.8 ± 20.0 |
| **Projected Climate (2040 – 2050)** | | | | | | |
| Avg. winter temp. (°F) | 37.1 ± 5.1<br>36.0 ± 5.5 | 33.2 ± 6.0<br>31.4 ± 6.8 | 37.1 ± 8.3<br>35.0 ± 8.6 | 47.1 ± 8.9<br>45.7 ± 9.2 | 50.2 ± 6.8<br>50.0 ± 7.1 | 48.2 ± 7.2<br>48.6 ± 7.5 |
| Avg. spring precip. (mm) | 4.9 ± 1.6<br>4.6 ± 1.4 | 4.8 ± 1.4<br>4.8 ± 1.5 | 3.0 ± 1.4<br>3.1 ± 1.3 | 1.8 ± 1.1<br>2.0 ± 1.1 | 3.1 ± 1.6<br>3.3 ± 1.5 | 4.4 ± 1.7<br>4.3 ± 1.5 |
| Hot, dry days (# days) | 5.6 ± 6.6<br>6.3 ± 6.4 | 6.1 ± 5.5<br>6.2 ± 5.0 | 1.9 ± 3.0<br>1.8 ± 3.1 | 5.9 ± 7.9<br>5.1 ± 6.3 | 31.0 ± 13.0<br>25.0 ± 12.7 | 18.1 ± 10.3<br>14.9 ± 8.6 |
| Cumulative temp (°F) | 20,740 ± 1,398<br>20,566 ± 1,433 | 20,464 ± 1,382<br>20,067 ± 1,462 | 18,690 ± 1,913<br>18,374 ± 1,996 | 20,397 ± 2,526<br>21,486 ± 2,405 | 24,795 ± 2,057<br>24,381 ± 2,003 | 24,225 ± 1,784<br>23,966 ± 1,757 |
| Total annual precip. (mm) | 1,418 ± 277<br>1,343 ± 279 | 1,193 ± 200<br>1,253 ± 216 | 1,406 ± 680<br>1,436 ± 670 | 1,354 ± 568<br>1,379 ± 618 | 915 ± 271<br>936 ± 291 | 1,409 ± 267<br>1,487 ± 333 |
| Temp. var. ($°F^2$) | 242.9 ± 41.4<br>261.3 ± 44.8 | 307.7 ± 54.8<br>334.0 ± 70.6 | 176.6 ± 63.3<br>187.5 ± 70.1 | 153.6 ± 57.6<br>152.3 ± 53.4 | 222.1 ± 56.8<br>199.7 ± 50.4 | 209.5 ± 56.4<br>196.5 ± 52.4 |
| Precip. var. ($mm^2$) | 53.6 ± 18.3<br>47.5 ± 16.1 | 45.3 ± 13.6<br>47.7 ± 15.6 | 36.8 ± 29.1<br>38.9 ± 29.7 | 64.5 ± 46.2<br>63.6 ± 46.3 | 41.9 ± 23.6<br>40.4 ± 21.0 | 65.7 ± 250<br>66.4 ± 26.7 |
| **Projected Climate (2090 – 2100)** | | | | | | |
| Avg. winter temp. (°F) | 41.2 ± 4.9<br>41.1 ± 4.9 | 38.5 ± 5.5<br>38.4 ± 5.5 | 40.1 ± 8.0<br>40.3 ± 8.0 | 50.1 ± 8.5<br>50.1 ± 8.5 | 54.7 ± 6.6<br>54.7 ± 6.6 | 53.0 ± 7.1<br>52.9 ± 7.1 |
| Avg. spring precip. (mm) | 3.7 ± 1.6<br>3.7 ± 1.5 | 4.6 ± 1.8<br>4.7 ± 1.9 | 2.9 ± 1.2<br>2.8 ± 1.1 | 1.6 ± 0.9<br>1.7 ± 0.9 | 2.9 ± 1.9<br>3.0 ± 1.9 | 3.6 ± 1.5<br>3.7 ± 1.6 |
| Hot, dry days (# days) | 16.9 ± 10.2<br>16.6 ± 10.3 | 20.2 ± 9.0<br>19.9 ± 9.1 | 5.8 ± 6.0<br>5.8 ± 5.9 | 12.6 ± 10.1<br>12.3 ± 9.5 | 37.0 ± 14.4<br>35.9 ± 14.1 | 28.2 ± 9.0<br>27.9 ± 9.0 |
| Cumulative temp (°F) | 22,234 ± 1,396<br>20,881 ± 1,407 | 22,078 ± 1,352<br>20,573 ± 1,386 | 19,936 ± 1,785<br>18,748 ± 1,948 | 23,048 ± 2,271<br>21,885 ± 2,368 | 26,174 ± 1,828<br>24,853 ± 2,036 | 25,632 ± 1646<br>24,349 ± 1710 |

| | | | | | | |
|---|---|---|---|---|---|---|
| Total annual precip. (mm) | 1,314 ± 301<br>1,368 ± 307 | 1,231 ± 262<br>1,266 ± 236 | 1,475 ± 713<br>1,403 ± 689 | 1,255 ± 493<br>1,370 ± 585 | 907 ± 337<br>952 ± 307 | 1,414 ± 274<br>1,459 ± 295 |
| Temp. var. (°F$^2$) | 264.5 ± 39.9<br>258.4 ± 37.0 | 322.1 ± 52.3<br>322.6 ± 53.6 | 185.8 ± 70.4<br>174.1 ± 69.3 | 155.6 ± 60.0<br>139.8 ± 51.5 | 215.0 ± 64.0<br>226.7 ± 53.3 | 206.3 ± 54.4<br>214.7 ± 54.6 |
| Precip. var. (mm$^2$) | 52.0 ± 17.6<br>51.5 ± 17.2 | 55.2 ± 20.4<br>49.9 ± 15.9 | 45.2 ± 36.5<br>37.5 ± 30.1 | 64.7 ± 40.9<br>67.0 ± 46.6 | 43.5 ± 27.2<br>43.2 ± 23.9 | 73.4 ± 25.4<br>67.6 ± 24.8 |

**Table S2.** Summary statistics for each region, including the population size (as of 2017), the number of counties, and the observed average climate conditions (mean ± standard deviation) from 2000 - 2017 for all US regions. Regional boundaries correspond to US Fish & Wildlife Service regions.

| | **Northeast** | **Midwest** | **Pacific** | **Pacific Southwest** | **Southwest** | **Southeast** |
|---|---|---|---|---|---|---|
| Pop. size | 74,464,546 | 61,721,093 | 13,265,462 | 42,534,692 | 41,339,800 | 73,429,199 |
| No. counties | 435 | 738 | 119 | 75 | 379 | 875 |
| Avg. winter temp. (°F) | 31.6 ± 6.3 | 25.6 ± 7.5 | 32.2 ± 6.8 | 42.3 ± 8.6 | 46.2 ± 7.5 | 44.8 ± 7.4 |
| Avg. spring precip. (mm) | 3.4 ± 1.4 | 3.6 ±1.6 | 1.6 ± 1.1 | 0.5 ± 0.7 | 2.7 ± 2.5 | 3.5 ± 1.7 |
| Hot, dry days (# days) | 4.5 ± 5.5 | 7.7 ± 7.8 | 4.2 ± 7.0 | 15.6 ± 18.0 | 16.7 ± 9.8 | 34.8 ± 14.4 |
| Cumulative temp (°F) | 19,169.9 ± 1,696.3 | 18,662.6 ± 1,746.5 | 17,637.9 ± 1,475.2 | 20,710.2 ± 2,465.4 | 24,018.9 ± 1,855.6 | 23,090.1 ± 1,731.4 |
| Cumulative precip. (mm) | 1,095.7 ± 253.7 | 935.5 ± 245.9 | 844.1 ± 624.0 | 517.7 ± 411.9 | 736.7 ± 434.3 | 1,238.1 ± 306.9 |
| Temp. variance (°F$^2$) | 302.8 ± 48.1 | 414.8 ± 89.0 | 225.7 ± 105.7 | 191.3 ± 83.6 | 266.1 ± 75.0 | 245.8 ± 64.6 |
| Precip. variance (mm$^2$) | 54.0 ± 26.6 | 44.4 ± 22.3 | 33.7 ± 47.5 | 30.5 ± 38.1 | 43.5 ± 37.5 | 74.7 ± 35.7 |

**Table S3.** Effect of climate and non-climate variables on Lyme disease incidence by region generated under the main model specification and using standard errors (SE) clustered at the county level. Only variables included in the best model, as determined by variable selection, are shown. The scaled coefficient estimates (Coef.) shown here reflect the standard deviation change in Lyme disease incidence for a one standard deviation change in the climate variable. The coefficients are scaled so that the effects of different variables are directly comparable. Statistically significant ($p < 0.05$) coefficients are denoted with *.

| | Northeast | | Midwest | | Pacific | | Pacific Southwest | | Southwest | | Southeast | |
|---|---|---|---|---|---|---|---|---|---|---|---|---|
| **Variable** | **Coef.** | **SE** | **Coef.** | **SE** | **Coef.** | **SE** | **Coef.** | **SE** | **Coef.** | **SE** | **Coef.** | **SE** |
| Avg. winter temp. | | | -0.073 | 0.116 | -0.967 | 0.982 | 0.119 | 0.296 | | | | |
| Avg. winter temp.$^2$ | | | 0.381* | 0.119 | 1.268 | 0.830 | 0.391 | 0.439 | | | | |
| Avg. spring precip. | 0.067 | 0.081 | -0.051* | 0.023 | | | 0.089 | 0.119 | -0.998 | 0.864 | | |
| Avg. spring precip.$^2$ | -0.094 | 0.055 | | | | | | | | | | |
| Hot, dry days | -0.302* | 0.074 | -0.264* | 0.055 | | | | | 0.151 | 0.123 | -0.029 | 0.020 |
| Hot, dry days $^2$ | 0.106* | 0.038 | 0.121* | 0.032 | | | | | | | | |
| Cumulative temp. | 1.034* | 0.225 | | | | | | | 1.589 | 1.085 | 1.928 | 1.359 |
| Cumulative temp.$^2$ | | | | | | | | | -2.127 | 1.292 | -2.405 | 1.462 |
| Total annual precip. | -0.141 | 0.153 | -0.046 | 0.082 | | | | | 1.192 | 1.021 | | |
| Total annual precip.$^2$ | 0.183 | 0.125 | -0.010 | 0.055 | | | | | | | | |
| Temp. variability | 0.365 | 0.378 | | | | | 0.112 | 0.999 | | | 0.813* | 0.253 |
| Temp. variability $^2$ | 0.131 | 0.344 | | | | | 0.224 | 0.430 | | | -0.473* | 0.207 |
| Precip. variability | | | 0.040 | 0.038 | | | | | -0.220 | 0.192 | | |
| Precip. variability $^2$ | | | 0.012 | 0.018 | | | | | | | | |
| Lag 'ticks' search | 0.168* | 0.044 | 0.016 | 0.015 | 0.014 | 0.039 | 0.049 | 0.080 | 0.020 | 0.048 | -0.016 | 0.014 |
| Poverty | -0.055 | 0.076 | 0.046 | 0.067 | | | | | 0.210 | 0.164 | | |
| Percent insured | | | | | | | | | | | -0.009 | 0.032 |
| Forest cover | 1.988 | 1.364 | -3.966 | 2.467 | -1.515* | 1.028 | -0.365 | 0.604 | | | 0.663 | 0.350 |
| Mixed dev. cover | | | | | | | | | 1.447 | 1.241 | 1.441* | 0.565 |
| $R^2$ | 0.728 | | 0.829 | | 0.405 | | 0.327 | | 0.309 | | 0.330 | |

**Table S4.** Effect of climate and non-climate variables on Lyme disease incidence by region, under model specification two in which only linear versions of climate predictors were used. Only variables included in the best model, as determined by variable selection, are shown. The scaled coefficient estimates (Coef.) shown here reflect the standard deviation change in Lyme disease incidence for a one standard deviation change in the climate variable. The coefficients are scaled so that the effects of different variables are directly comparable. The standard errors (SE) shown are clustered by the agricultural statistics district (see Methods: Statistical analysis). Statistically significant ($p < 0.05$) coefficients are denoted with *.

| | **Northeast** | | **Midwest** | | **Pacific** | | **Pacific Southwest** | | **Southwest** | | **Southeast** | |
|---|---|---|---|---|---|---|---|---|---|---|---|---|
| **Variable** | **Coef.** | **SE** | **Coef.** | **SE** | **Coef.** | **SE** | **Coef.** | **SE** | **Coef.** | **SE** | **Coef.** | **SE** |
| Avg. winter temp. | | | 0.277* | 0.136 | 0.223 | 0.447 | 0.337 | 0.221 | | | | |
| Avg. spring precip. | -0.053 | 0.038 | -0.067 | 0.051 | | | 0.105 | 0.080 | -1.027 | 0.849 | | |
| Hot, dry days | -0.193* | 0.064 | -0.154* | 0.050 | | | | | 0.153 | 0.138 | | |
| Cumulative temp | 0.965* | 0.481 | | | | | | | -0.552 | 0.688 | -0.464* | 0.231 |
| Total annual precip. | | | | | | | | | 1.215 | 0.992 | | |
| Temp. variability | 0.480* | 0.151 | | | | | 0.452 | 0.249 | | | 0.276* | 0.101 |
| Precip. variability | 0.057 | 0.032 | | | | | | | -0.224 | 0.178 | | |
| Lag 'ticks' search | 0.170* | 0.074 | 0.017 | 0.017 | -0.005 | 0.045 | 0.057 | 0.062 | 0.010 | 0.067 | -0.026 | 0.020 |
| Poverty | -0.052 | 0.090 | 0.032 | 0.073 | | | -0.075 | 0.067 | 0.207 | 0.133 | | |
| Percent insured | | | | | | | | | | | -0.001 | 0.038 |
| Forest cover | 1.822 | 1.311 | -3.850 | 3.721 | -1.063 | 0.750 | -0.458 | 0.434 | | | 0.780 | 0.416 |
| Mixed dev. cover | | | | | | | | | 1.401 | 1.619 | 1.383* | 0.686 |
| $R^2$ | 0.726 | | 0.827 | | 0.401 | | 0.325 | | 0.309 | | 0.326 | |
| **Model with only climate and dummy variables** | | | | | | | | | | | | |
| $R^2$ | 0.680 | | 0.766 | | 0.230 | | 0.137 | | 0.111 | | 0.145 | |
| **Model with only non-climate and dummy variables** | | | | | | | | | | | | |
| $R^2$ | 0.712 | | 0.820 | | 0.400 | | 0.310 | | 0.258 | | 0.320 | |
| **Model with only county dummy variable** | | | | | | | | | | | | |
| $R^2$ | 0.606 | | 0.700 | | 0.156 | | 0.114 | | 0.090 | | 0.149 | |
| **Model with only year dummy variable** | | | | | | | | | | | | |
| $R^2$ | 0.045 | | 0.018 | | 0.028 | | 0.014 | | 0.007 | | 0.010 | |

**Table S5.** Projected change in the number of Lyme disease cases, relative to hindcasted 2010 – 2020 levels, made using model specification two in which only linear versions of climate variables were used (see Methods: Model validation). Point estimates and 95% prediction intervals are shown.

| | **Upper climate change scenario (RCP8.5)** | | **Moderate climate change scenario (RCP4.5)** | |
|---|---|---|---|---|
| | 2040 - 2050 | 2090 - 2100 | 2040 – 2050 | 2090 - 2100 |
| Northeast | 21,467 [113, 42,820] | 42,538 [18,409, 66,666] | 6,972 [-14,890, 28,883] | 4,014 [-20,888, 28,918] |
| Midwest | -2,784 [-11,111, 5,544] | -6,011 [-14,878, 2,856] | 1,725 [-5,703, 9,152] | 2,926 [-4,740, 10,593] |
| Pacific | 57 [-207, 320] | 130 [-345, 604] | 8 [-218, 238] | 56 [-292, 404] |
| Pacific Southwest | -49 [-1,796, 1,698] | -174 [-2,156, 1,808] | 30 [-1,635, 1,695] | -81 [-1,953, 1,791] |
| Southwest | -71 [-1,228, 1,086] | -313 [-2,009, 1,383] | 63 [-1,107, 1,233] | 135 [-1,589, 1,859] |
| Southeast | 1,522 [310, 2,735] | 3,640 [1,904, 5,374] | 257 [-914, 1,427] | 341 [-1,126, 1,808] |
| US Total | 20,142 [-13,919, 59,663] | 39,810 [-2,675, 78,691] | 9,055 [-24,467, 42,628] | 7,391 [-30,588, 45,373] |

**Table S6.** Projected change in the number of Lyme disease cases, relative to hindcasted 2010 – 2020 levels, assuming no change in county population sizes from 2017 (see Methods: Lyme disease projections). Point estimates and 95% prediction intervals are shown.

| | **Upper climate change scenario (RCP8.5)** | | **Moderate climate change scenario (RCP4.5)** | |
|---|---|---|---|---|
| | 2040 - 2050 | 2090 - 2100 | 2040 – 2050 | 2090 - 2100 |
| Northeast | 18,885<br>[-624, 38,395] | 40,320<br>[18,434, 62,206] | 6,247<br>[-13,899, 26,393] | -3,352<br>[-29,194, 22,091] |
| Midwest | -2,454<br>[-10,561, 5,652] | -4,460<br>[-14,352, 5,433] | 2,373<br>[-5,499, 10,244] | 1,116<br>[-10,072, 12,304] |
| Pacific | 18<br>[-177, 212] | 34<br>[-245, 312] | -5<br>[-172, 161] | 14<br>[-187, 214] |
| Pacific Southwest | -92<br>[-1,560, 1,376] | -220<br>[-1,878, 1,438] | -132<br>[-1,448, 1,185] | -214<br>[-1,604, 1,177] |
| Southwest | -135<br>[-908, 638] | -343<br>[-1,214, 528] | -120<br>[-887, 646] | -169<br>[-953, 615] |
| Southeast | 637<br>[-310, 1,583] | 962<br>[-213, 2,137] | 135<br>[-798, 1,067] | 278<br>[-733, 1,289] |
| US Total | 16,859<br>[-14,140, 47,856] | 36,293<br>[532, 72,054] | 8,498<br>[-22,703, 39,696] | -2,327<br>[-42,743, 37,690] |

**Table S7**. Population sizes by region. For 2017, population estimates are from the United States Census Bureau (USCB). For 2050 and 2100, population projections (under SSP2) are from Hauer et al. 2019.

| | **2017** | **2050** | **2100** |
|---|---|---|---|
| Northeast | 74,464,546 | 81,948,939 | 83,080,395 |
| Midwest | 61,721,093 | 63,835,338 | 59,824,981 |
| Pacific | 13,265,462 | 18,067,864 | 22,779,701 |
| Pacific Southwest | 42,534,692 | 53,832,936 | 57,299,030 |
| Southwest | 41,339,800 | 62,932,217 | 86,639,719 |
| Southeast | 73,429,199 | 94,766,152 | 111,891,023 |
